# Enriched zones of embedded ribonucleotides are associated with DNA replication and coding sequences in the human mitochondrial genome

**DOI:** 10.1101/2023.04.05.535745

**Authors:** Penghao Xu, Taehwan Yang, Deepali L. Kundnani, Mo Sun, Stefania Marsili, Alli L. Gombolay, Youngkyu Jeon, Gary Newnam, Sathya Balachander, Veronica Bazzani, Umberto Baccarani, Vivian S. Park, Sijia Tao, Adriana Lori, Raymond F. Schinazi, Baek Kim, Zachary F. Pursell, Gianluca Tell, Carlo Vascotto, Francesca Storici

**Affiliations:** School of Biological Sciences, Georgia Institute of Technology, Atlanta, GA, USA; Department of Medicine, University of Udine, Udine, Italy; General Surgery Clinic and Liver Transplant Center, University-Hospital of Udine; Department of Biochemistry and Molecular Biology, Tulane Cancer Center, Tulane University of Medicine, New Orleans, Louisiana; Department of Pediatrics, School of Medicine, Emory University, Atlanta, GA, USA; Department of Psychiatry and Behavioral Sciences, Emory University, Atlanta, Georgia, USA; Department of Population Science, American Cancer Society, Atlanta, GA; Laboratory of Molecular Biology and DNA repair, Department of Medicine, University of Udine, Udine, Italy; IMol Polish Academy of Sciences, Warsaw, Poland

**Keywords:** Ribonucleotides, human mitochondrial DNA, Ribose-Map, DNA replication, ribose-seq, yeast mitochondrial DNA, hepatocellular carcinoma, human embryonic kidney HEK293T cells, human embryonic stem cells hESC-H9, lymphocytes CD4^+^T.

## Abstract

Abundant ribonucleoside triphosphate (rNTP) incorporation in DNA by DNA polymerases in the form of ribonucleoside monophosphates (rNMPs) is a widespread phenomenon in nature, resulting in DNA structural change and genome instability. The rNMP distribution, characteristics, hotspots, and association with DNA metabolic processes in human mitochondrial DNA (hmtDNA) remain mostly unknown. Here, we utilize the *ribose-seq* technique to capture embedded rNMPs in mtDNA of six different human cell types with wild-type or mutant ribonuclease (RNase) H2 genotype. The rNMPs are preferentially embedded in the DNA of the light strand in most cell types studied, but not in the liver-tissue cells, in which the rNMPs are dominant on the heavy strand of hmtDNA. We uncover common rNMP hotspots and conserved rNMP-enriched zones across the entire hmtDNA, including in the replication-control region, which may result in the suppression of mtDNA replication. We also show that longer coding sequences have a significantly higher rNMP-embedment frequency per nucleotide. While the composition of the embedded rNMPs varies among the different cell types, by studying the genomic context of embedded rNMPs, we detected common rNMP-embedment patterns in hmtDNA. The genomic contexts of rNMPs found in hmtDNA are mainly distinct from those found in yeast mtDNA, highlighting a unique signature of rNTP incorporation by hmtDNA polymerase γ.

## Background

Human mitochondrial DNA (hmtDNA) consists of two strands and 37 genes, including 13 coding sequences (CDSs), 2 ribosomal RNA (rRNA) genes, and 22 transfer RNA (tRNA) genes (1). Maintaining the mitochondrial DNA (mtDNA) functions is essential to human cells. Malfunction of the hmtDNA-replication machinery leads to mtDNA-depletion syndromes or mtDNA-deletion disorders, resulting in respiratory-chain deficiency and neuromuscular symptoms in patients (2). Ribonucleoside triphosphates (rNTPs) are the basic building blocks of RNA. Due to the structural similarity to deoxyribonucleoside triphosphates (dNTPs), rNTPs are abundantly incorporated into DNA in the form of ribonucleoside monophosphates (rNMPs) by DNA polymerases despite their ability to differentiate the sugar moieties (3–5). The incorporated rNMPs constitute the most abundant non-standard nucleotides (nt) in the DNA of eukaryotic and prokaryotic genomes (6–9). The rNMP-embedment rates of replicative DNA polymerases, including DNA Pol α, δ, ε for eukaryotic nuclear DNA, and Pol γ for mtDNA have been studied *in vitro* (7, 10–12). At normal dNTP concentrations, human Pol γ incorporates approximately one rNMP per ∼2,300 nt (12, 13). Majority of the rNMPs embedded in nuclear DNA are efficiently removed through the ribonucleotide-excision repair (RER) pathway, which is initiated by ribonuclease (RNase) H2 (14). Inactivation of RER can be caused by mutations in any of the of three RNase H2 subunits (15). Deficiency in rNMP removal from nuclear DNA is associated with human diseases such as Aicardi-Goutières Syndrome (AGS), Systemic Lupus Erythematosus (SLE), and different types of cancer (7, 16–18). In yeast and human mtDNA, the embedded rNMPs cannot be repaired by RER due to the absence of RNase H2 in the organelle (19–21). As a result, hmtDNA has abundant rNMPs (21, 22). Although human Pol γ can efficiently bypass single rNMPs with high fidelity (13), the presence of rNMPs in DNA can lead to DNA fragility and structural change, and affect DNA-protein interactions (23, 24).

Only recently, different rNMP-mapping techniques have been developed including ribose-seq (22), emRiboSeq (25), HydEn-seq (26), Pu-seq (27), RiSQ-seq (28), and GLOE-Seq (29). These techniques can efficiently capture the embedded rNMPs in genomic DNA and facilitate genomic studies on rNMPs. Bioinformatic tools have also been developed to map the rNMPs in genomic DNA (30), optimize the rNMP capture techniques (31), and analyze rNMP embedment characteristics (32, 33). Many of the rNMP-embedment characteristics in mtDNA of the yeast *Saccharomyces cerevisiae*, *Saccharomyces paradoxus*, *Schizosaccharomyces pombe*, and the green alga *Chlamydomonas reinhardtii* have been recently illustrated in detail (21, 34). However, current knowledge about rNMP embedment in hmtDNA has been limited to count and composition in HeLa and fibroblast cells (12). It remains unknown whether the rNMPs found in hmtDNA have a biased distribution, whether rNMP-hotspots and patterns exist in hmtDNA, and whether these rNMP features differ in different human cell types. Therefore, the characteristics and patterns of rNMPs found in hmtDNA in different cell types, and their relation to hmtDNA genes and replication still need to be investigated. Uncovering the specific features of rNMPs found in hmtDNA can reveal the relation between rNMPs in DNA and many human diseases associated with the alteration of mtDNA metabolism and stability.

In this study, we analyze 32 hmtDNA libraries of six different human cell types, which we constructed using the ribose-seq technique. Our results identify strand bias, hotspots, and preferred patterns of rNMPs in hmtDNA. We uncover the presence of rNMP-enriched zones (REZs), including within the replication control region, and we discuss their potential effects on hmtDNA replication. Our detailed analysis of embedded rNMPs in hmtDNA reveals specific rNMP-embedment characteristics and an unprecedented association between rNMP-embedment frequency and gene size. In summary, we identify unique characteristics of embedded rNMPs in hmtDNA and unveil an unexpected relationship among the embedded rNMPs, mtDNA replication, and coding sequences in the hmtDNA genome.

## Results

### rNMP-embedment preference on the light strand of hmtDNA

The two strands of hmtDNA are named the light strand and the heavy strand, due to their differences in molecular weight. Twelve of the 13 CDSs are located on the light strand and are transcribed as one mRNA from the heavy strand promoter (HSP) (35). Therefore, the heavy strand mainly works as a template strand for the transcription of mRNA. In this study, we constructed 32 ribose-seq libraries from mtDNA of six different human cell types: lymphocytes CD4^+^T, embryonic stem cells hESC-H9, cells derived from distal or tumor liver-tissue biopsies (DLTB and TLTB), whole blood cells from a patient affected by post-traumatic stress disorder (PTSD) and control (WB-GTP PTSD and WB-GTP control), colorectal carcinoma cell line HCT116, and human embryonic kidney cells (HEK293T), having either wild-type ribonuclease (RNase) H2, or having the catalytic subunit of RNase H2 knocked-out (RNH2A-KO) in two different cell clones (RNH2A-KO T3-8 and RNH2A-KO T3-17) (**Additional file 1**, and see Methods). These cells are classified in our study into 10 different cell categories: CD4^+^T, hESC-H9, DLTB, TLTB, WB-GTP control, WB-GTP PTSD, HCT116, HEK293T, RNH2A-KO T3-8, and RNH2A-KO T3-17. In the preparation of the ribose-seq libraries, we utilized different fragmentation methods to cut hmtDNA, including dsDNA fragmentase, three distinct restriction enzyme (RE) combinations optimized using the RESCOT software (31), and dsDNA fragmentase together with RE combinations. With the paired-end sequencing reads, we adapted the Ribose-Map protocol (30, 36) to align the rNMPs to the circular mtDNA of *Homo sapiens* GRCh38 reference genome and recovered the unaligned rNMPs at the beginning and end in the reference sequence (see Methods). The library information including cell types, RNase H2 genotypes, fragmentation methods, and count of rNMPs is presented in **Additional file 1**. By analyzing the data from the 32 ribose-seq libraries, we found that the rNMP embedment in hmtDNA of the six cell types studied is biased between the light and the heavy strands. Specifically, rNMPs are preferentially embedded on the light strand, which is the non-template strand relative to transcription of the long transcript generated from the HSP, in most of the ribose-seq libraries of the cell types examined. In the libraries derived from HEK293T cells that are wild-type or knocked out for RNase H2A, the bias is less prominent or not significant. Interestingly, the ribose-seq libraries derived from liver-tissue samples (DLTB and TLTB) show a dominant presence of rNMPs on the heavy strand, or template strand of hmtDNA **(Fig. 1A)**. This biased-rNMP presence on the light or heavy strand of hmtDNA is not due to the uneven abundance of the heavy or light DNA strands in the cells, because the DNA-seq results of mtDNA extracted from each of the cell types show a balanced count of reads aligned to the light and heavy strands (**Additional file 2**).

**Figure 1.**
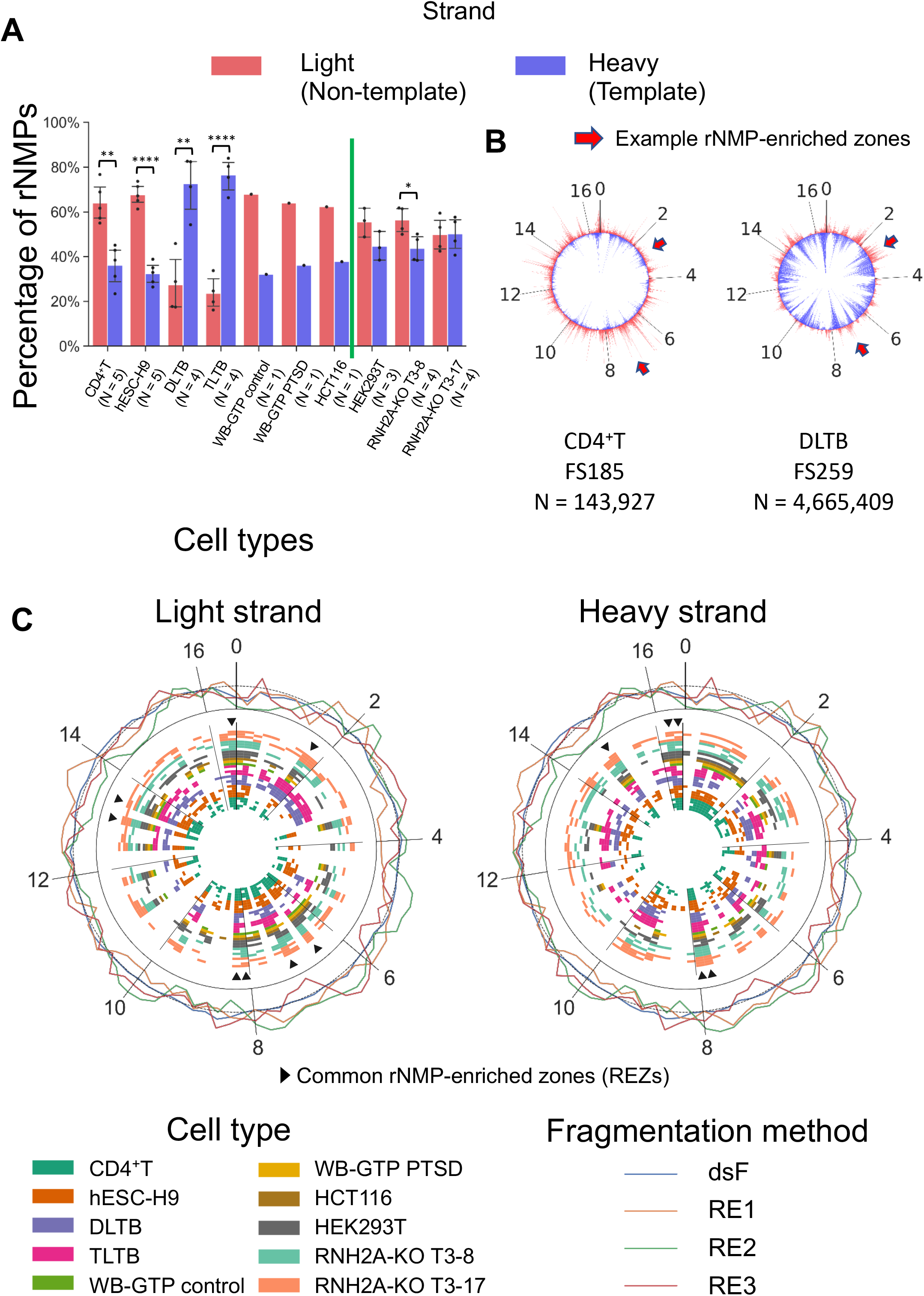
Biased rNMP distribution in hmtDNA. (**A**) Bar graph showing the percentage of rNMP embedment on the light (red bars) and heavy (blue bars) strands in hmtDNA. The cell type is indicated below each bar. The green vertical line separates cell types with RNH2A WT and KO. Two-tailed Mann-Whitney *U* test is performed to check the significance of the light/heavy strand bias for all cell types with N >= 3. *: 0.01 < P < 0.05; **: 0.001 < P < 0.01; ***: 0.0001 < P < 0.001; ****: 0.00001 < P < 0.0001. **(B)** rNMP frequency at each nucleotide on the light (outer-circle red spikes) and heavy (inner-circle blue spikes) strands of hmtDNA. Examples of consistent enrichment of rNMPs on the light strand are indicated by red arrows. **(C)** The rNMP-enriched zones (REZs) on the light and heavy strands of hmtDNA. Each circle represents one library. Each block in each circle represents a REZ, which is a 200-nt zone having an rNMP enrichment factor > 1. The color of the circles represents the cell type and genotype. The common REZs are 200-nt zones that are REZs in all cell categories and genotypes, and more than 80% of the libraries studied. The common REZs are indicated by the triangle markers and listed in **Additional file 4**. The outer circle shows the background coverage of RE and dsDNA fragmentase. The mean background coverage of the whole hmtDNA (Enrichment factor = 1) is represented as the dotted line in the outer circle.

As a comparison, we performed the same strand-distribution analysis of embedded rNMPs in budding yeast mtDNA. We analyzed 11 ribose-seq libraries derived from wild-type RNase H2 strains of *S. cerevisiae* and 8 libraries derived from mutant-RNase H2 strains with the catalytic subunit deleted (*rnh201-*null) that were built in a previous study (21). These yeast libraries derive from six different strains and have been fragmented using three different RE combinations. The *S. cerevisiae* mtDNA contains 35 genes, including 8 CDS, 1 ribosomal protein, 2 rRNA genes, and 24 tRNA genes (37). All yeast mtDNA CDSs are located on the forward strand. By analyzing these yeast libraries, we found that the rNMPs are preferentially embedded on the reverse strand in both wild-type and *rnh201*-null libraries **(Additional file 3: Fig. S1A)**. Considering that the reverse strand is the template strand of all CDSs in yeast mtDNA during the process of transcription, yeast mtDNA has an opposite strand bias of rNMP distribution to that found in hmtDNA relative to transcription in most of the human cell types studied here, but a similar bias to that found in hmtDNA libraries derived from the liver-tissue cells.

### rNMP-enriched zones and rNMP hotspots in hmtDNA

To better understand the rNMP-embedment characteristics in hmtDNA, we plotted the rNMP distribution maps at the single-nucleotide level **(Fig. 1B)** and **Additional file 3: Fig. S2)**. From the distribution maps, we found that the rNMPs are distributed unevenly along either strand of the mtDNA. We identified some genomic zones with enrichment of incorporated rNMPs, which have higher rNMP-embedment frequency relative to the hmtDNA average. We termed these zones as rNMP-enriched zones (REZs). We calculated the rNMP-enrichment factor of all 166 bins of 200-nt size covering both mtDNA strands for all the rNMP libraries within the 10 cell categories listed above and we identified the REZs. Surprisingly, we found that the distribution of REZs in the different libraries shows high consistency (**Fig. 1C**). We revealed eight common REZs on the light strand and five common REZs on the heavy strand which are present in all 10 cell categories and more than 80% of the libraries (**Fig. 1C, Additional file 4**). The presence of these common REZs is therefore independent from the cell type. To check if the REZs represent an artificial result caused by the different methods of hmtDNA fragmentation, we performed DNA-seq and calculated the read coverage in CD4^+^T libraries fragmented by each of the three RE combinations, and by dsDNA Fragmentase (outer circle in **Fig. 1C**). Results show that the libraries constructed using dsDNA Fragmentase keep a consistent coverage all over the mtDNA. The libraries prepared using restriction enzymes (REs) have some fluctuation among different hmtDNA regions due to the different recognition patterns. However, the common REZs are not enriched in either high-read-coverage regions or low-read-coverage regions, suggesting that the common REZs are uncorrelated with the different fragmentation methods. Hence, the common REZs are intrinsic properties of rNMP embedment in hmtDNA.

To determine whether REZs are present in the yeast mtDNA, we generated the rNMP-distribution maps of previously published ribose-seq and emRiboSeq libraries of *S. cerevisiae* (21, 25). Similarly to what was found in hmtDMA, among all the yeast mtDNA ribose-seq libraries analyzed, common REZs existed independently from i) the RNase H2 genotype; ii) the RE set used for fragmenting the genome (RE1, RE2, or RE3); iii) the rNMP-mapping technique used, which is particularly relevant because emRiboSeq utilizes a protocol for genome fragmentation and capture of rNMPs different from ribose-seq (25) (**Additional file 3: Fig. S1B**). Overall, the results obtained for human and yeast mtDNA show an uneven distribution of rNMPs in the mtDNA genome and highlight the presence of preferred regions of rNMP embedment. The consistency of common REZs across the human and yeast mtDNAs that we studied may reflect specific metabolic activities or structural features of mtDNA.

The rNMP hotspots are single-nucleotide locations with one or more rNMPs in all of the 32 libraries studied. We calculated the average rNMP-embedment enrichment factor for libraries derived from the same cell type and found that the rNMP hotspots are present in regions of high rNMP-embedment frequency (**Fig. 2**). We identified common rNMP hotspots in hmtDNA in all libraries we analyzed (**Fig. 2A**), as well as unique rNMP hotspots for CD4^+^T, hESC-H9, DLTB, HEK293T, and HEK293T RNH2A-KO (RNH2A-KO T3-8 and RNH2A-KO T3-17) subsets (**Fig. 2B-F**, **Additional file 5**). The top 20 hotspots with the highest median enrichment factor are selected and marked in **Fig. 2**. Most selected rNMP hotspots (17 out of 20) in the combined-hmtDNA libraries are present on the light strand and inside the non-template strand of the coding sequences (**Fig. 2A**). Most of them are clustered together in two windows of ∼100-nt length located on the non-template strand of the NADH dehydrogenase 5 (MT-ND5) gene, and the Cytochrome C Oxidase subunit I (MT-CO1) gene. 70% of the common-rNMP hotspots are rCMPs, suggesting that rCMPs are strongly preferred as the rNMP hotspots in hmtDNA. Considering the individual subsets (CD4^+^T, DLTB, hESC-H9, HEK293T, and HEK293T RNH2A KO), we identified abundant rNMP hotspots and selected the top 20 (**Fig. 2B-F, Additional file 5**). The rNMP hotspots in each subset have different strand preferences that are in line with the strand-biased distribution of the rNMPs in the subset. For CD4^+^T, hESC-H9, and HEK293T libraries, there are more rNMP hotspots on the light strand **(Fig. 2B, C, E)**. On the contrary, in the DLTB libraries, we see rNMP hotspots mainly present on the heavy strand (**Fig. 2D**), which is consistent with the dominant presence of rNMPs on the heavy strand in these libraries (**Fig. 1A**). Furthermore, the clusters of rNMP hotspots also exist in the control region in DLTB libraries and on the template strand of Cytochrome B (MT-CYB) and Cytochrome C Oxidase II (MT-CO2) genes in the HEK293T RNH2A-KO cells **(Fig. 2D, F)**. Moreover, the rNMP hotspots in each subset show a strong composition preference. In the CD4^+^T and DLTB libraries, almost all rNMPs are rAMPs (CD4^+^T: 19 out of 20, **Fig. 2B**; DLTB: 20 out of 20, **Fig. 2D**). In the hESC-H9 libraries, almost all unique rNMP hotspots are rCMPs (19 out of 20, **Fig. 2C**). In the HEK293T libraries, all unique rNMP hotspots are rCMPs or rGMPs (**Fig. 2E**), and in the HEK293T RNH2A-KO libraries, most unique rNMP hotspots are rGMPs (14 out of 20, **Fig. 2F**). The consistency of the rNMP hotspots may play a vital role in hmtDNA gene function, especially in the clustering regions.

**Figure 2.**
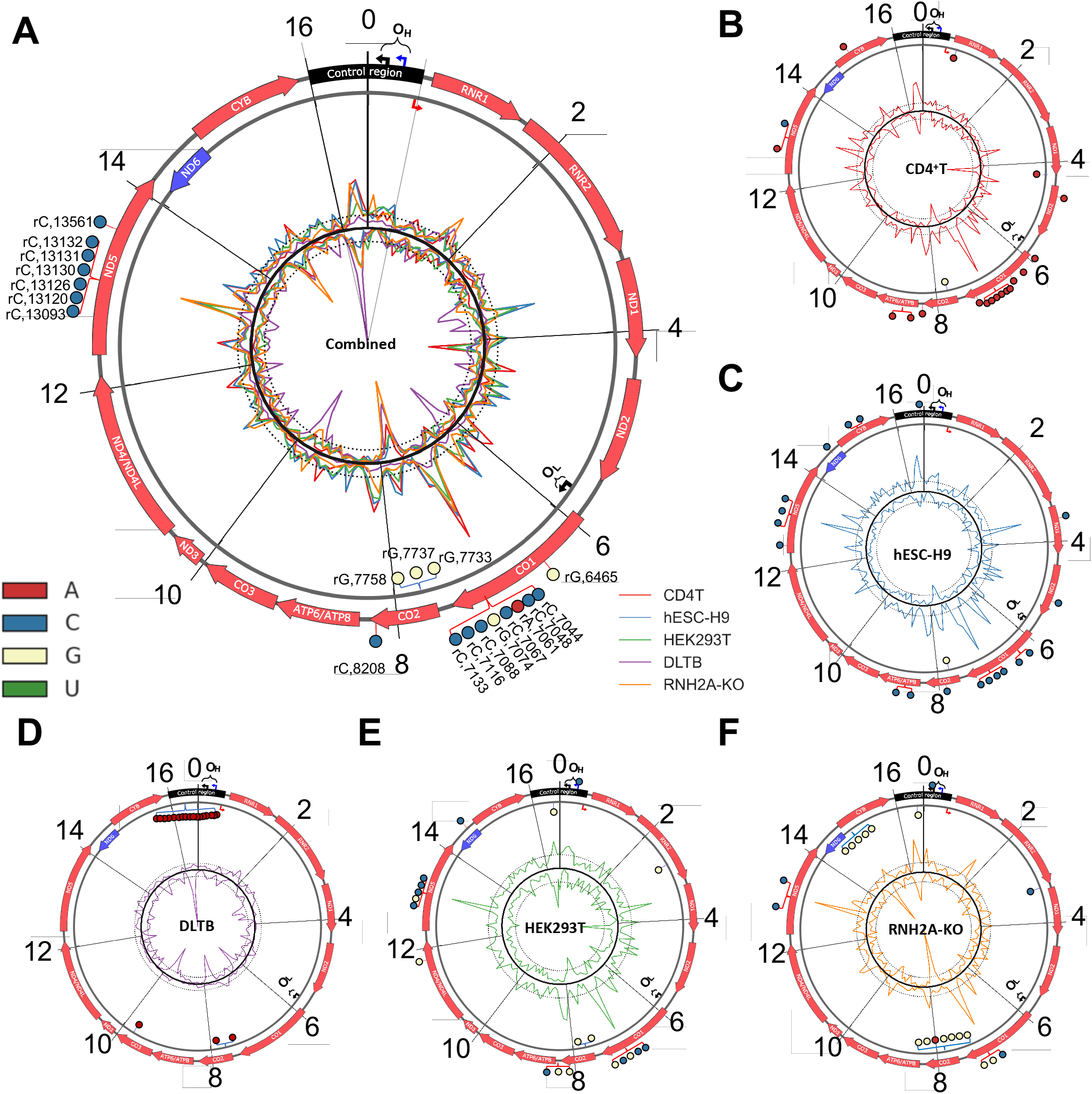
rNMP hotspots in hmtDNA.

Circular plots showing the rNMP-embedment frequency and rNMP hotspots in hmtDNA for **(A)** all hmtDNA libraries, **(B)** CD4T, **(C)** hESC-H9, **(D)** DLTB, **(E)** HEK293T RNH2A WT, and **(F)** HEK293T RNH2A KO libraries (Both RNH2A-KO T3-8 and RNH2A-KO T3-17). The coding sequences on the light and heavy strands are indicated as red and blue arrows on the outer circle. The control region is marked as black on the top. Two black braces represent the replication origin regions of the heavy strand (OriH, OH) and light strand (OriL, OL), and the black bent arrows show the actual replication start sites. The blue and red bent arrows indicate the heavy-strand promoter (HSP) and light-strand promoter (LSP), respectively. The outer and inner lines of the inner circle are the rNMP-enrichment factor of the light and heavy strands of cell categories containing more than 3 libraries. Two dotted lines around the circle indicate the mean value of rNMP-embedment frequency (Enrichment factor = 1. See **Methods**). The top 20 common hotspots in each subset and all hmtDNA on the light and heavy strands are marked as colored pins in the outer circle. The types of rNMP hotspots are shown in different colors. The type and location of all hotspots are also listed in **Additional file 5**.

### rNMP-embedment characteristics in the control region of hmtDNA

There is only one long non-coding region in the hmtDNA located around the zero position in the hmtDNA (black box in **Fig. 2)**, which is called the control region and contains essential genetic elements for mtDNA transcription and replication including the heavy-strand promoter (HSP), the light-strand promoter (LSP), and the termination-associated sequences (TAS) (**Fig. 3A**) (38). RNA polymerase POLRMT performs the transcription starting at HSP and LSP, and *in vitro* studies have shown that POLRMT can be terminated by embedded rNMPs in DNA (39). The heavy-strand replication origin (OriH, OH) and the D-loop, a special three-strand region containing a displaced light strand, a heavy strand, and a hmtDNA replication failure product 7S DNA are also present in the control region. We identified three strong common REZs in the control region, two on the light strand and one on the heavy strand, located in the D-loop region right before the OriH (**Fig. 3B, C**). These three REZs exist in all subsets and cell types we analyzed (**Additional file 3: Fig. S3**). The abundance of embedded rNMPs in these REZs is in line with the strand preference of the whole hmtDNA in the different cell types. For most cell types, there are more embedded rNMPs in the light-strand REZs. However, the liver-tissue libraries, DLTB and TLTB, have fewer rNMPs in the light-strand REZs and more rNMPs in the heavy-strand REZs (**Fig. 3B**). Abundant rNMP hotspots also clustered in the same zone on the heavy strand in DLTB libraries (**Fig. 2D**). Compared to the wild-type RNase H2 cells, similar and stronger REZs exist in HEK293T RNH2A-KO libraries (**Fig. 3C and Additional file 3: Fig. S3**). Moreover, we compared the rNMP embedment frequency in the control region to the published ChIP-seq data of DNA Pol γ and mtDNA helicase TWINKLE (40). We found that the ChIP-seq signal is lower in the REZs compared to the surrounding regions, which suggests that the binding activity of Pol γ and TWINKLE is lower in the REZs of both strands (**Additional file 3: Fig. S4**). The rNMPs in the D-loop region may inhibit the DNA binding activity of Pol γ and TWINKLE, suppressing hmtDNA replication.

**Figure 3.**
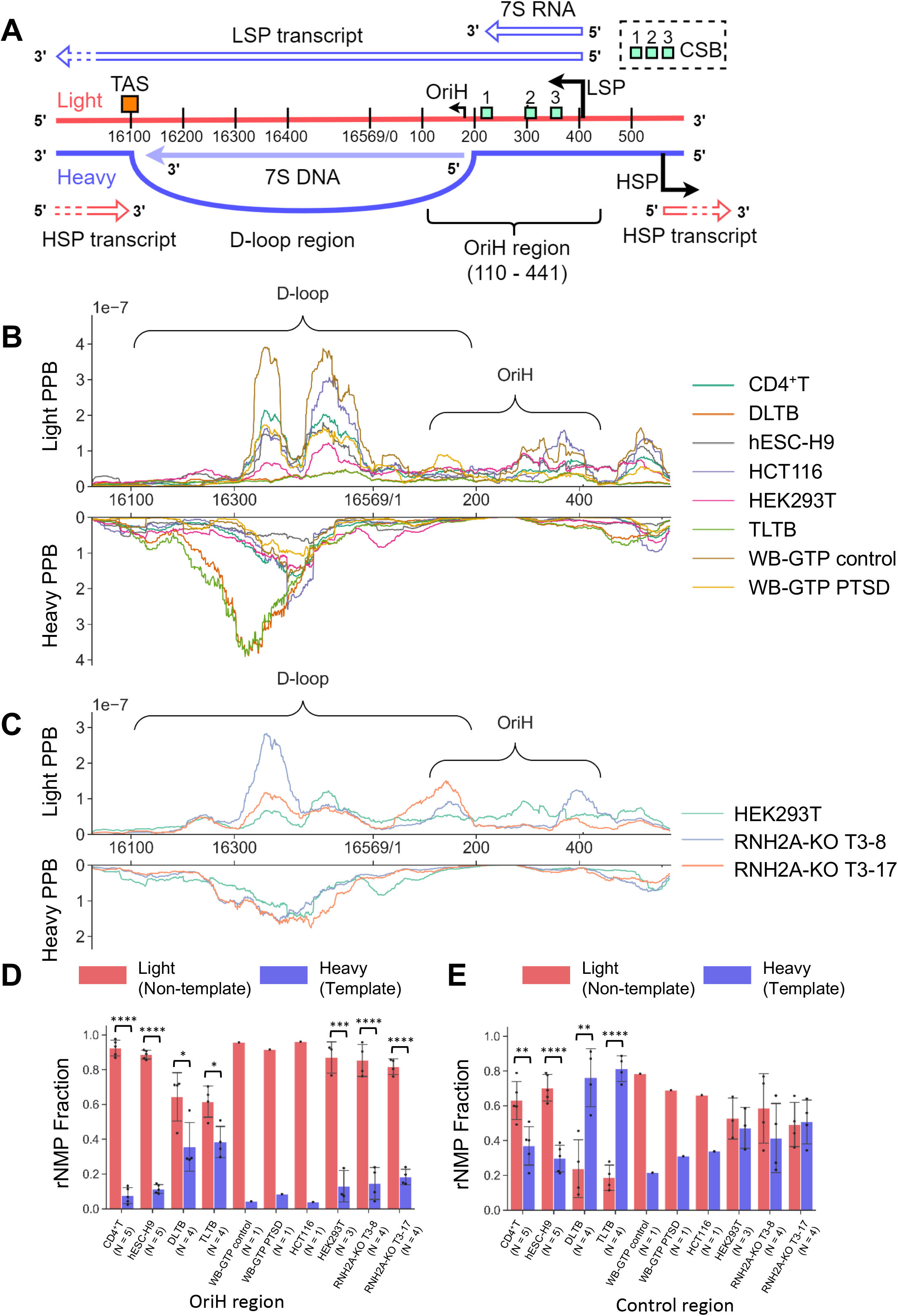
Conserved rNMP-enriched zones in the control region. (**A**) Scheme of genetic elements in the hmtDNA control region. **(B, C)** rNMP-embedment probability per base (PPB) in the hmtDNA-control region for (B) all RNH2A WT-cell types and **(C)** all HEK293T libraries with RNH2A KO or WT. The rNMP PPB on the light strand is on the upper part, while rNMP PPB on the heavy strand is on the bottom part. **(D, E)** The rNMP percentage on the light (red) and heavy (blue) strand for **(D)** OriH region and **(E)** control region in mtDNA. Two-tailed Mann-Whitney *U* test is performed to check the significance of the leading/lagging strand bias for all cell types with N >= 3. *: 0.01 < P < 0.05; **: 0.001 < P < 0.01; ***: 0.0001 < P < 0.001; ****: 0.00001 < P < 0.0001.

In hmtDNA, the light-strand transcripts start at LSP. Some of them are prematurely terminated and serve as primers for the heavy-strand replication (41, 42). The transition from terminated, priming RNAs to DNA happens at many different points around two conserved sequence blocks (CSB), CSB2 and CSB3 (43–46). Hence, the region containing these transition points has been known as the OriH region (1, 43) (**Fig 3A**). The parental-light strand, heavy strand, and the early-terminated transcript (7S RNA) form an R-loop in the OriH region (47). In this region, there are significantly more rNTPs incorporated on the light strand in all cell types (**Fig. 3D**). Even in the liver-tissue cells and HEK293T RNH2A-KO T3-17 libraries, where rNTPs are preferentially incorporated on the heavy strand in the whole hmtDNA, there are more rNTPs incorporated on the light strand in the OriH region. In contrast, the strong light-strand preference does not exist in the whole control region (**Fig. 3E**).

### Longer CDSs have a higher rNMP-embedment frequency on the non-template strand

We checked the rNMP-embedment frequency on both strands of all 13 CDSs in the hmtDNA. The rNMP-embedment frequency in the CDS region is in line with the strand preference of the whole mtDNA. In most cell types, there are abundant rNTPs incorporated on the light strand of DNA, which is the non-template strand for all CDSs except NADH dehydrogenase 6 (MT-ND6).

In the liver-tissue libraries, there are more rNTPs incorporated on the heavy strand, which is the template strand for all CDSs except MT-ND6 (**Fig. 4A and Additional file 3: Fig. S5A**). An unexpected finding is that the longer CDSs have more rNTPs incorporated on the non-template strand even after the normalization by CDS size. A significant positive correlation between CDS size and rNMP-embedment frequency is detected in all cell types despite the strand preference and RNase H2 genotype (Spearman’s r > 0.6, **Fig. 4B and Additional file 3: Fig. S5B, C**). To validate our finding, we generated three control groups with artificial rNTPs randomly incorporated in DNA. The results show that the CDS size is irrelevant to the rNMP-embedment frequency in the control group on the non-template strand (Spearman’s r < 0.2**, Additional file 3: Fig. S5A, B**). There is also no correlation between the CDS size and rNMP-embedment frequency on both template and non-template strands in yeast mtDNA (**Additional file 3: Fig. S6A, B**).

**Figure 4.**
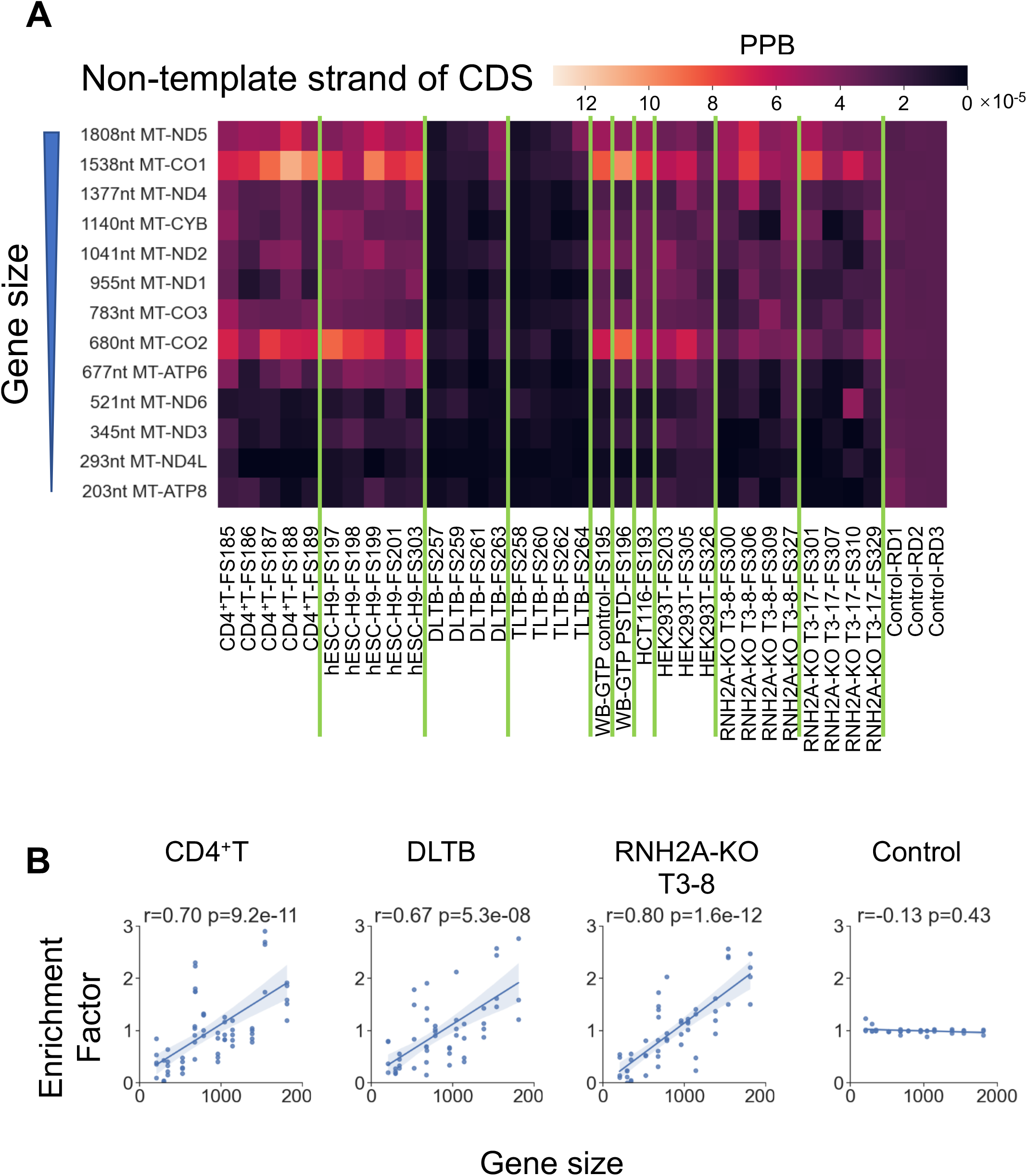
Longer CDSs have a higher rNMP-embedment frequency on the non-template strand. (**A**) rNMP-embedment probability per base (PPB) on the non-template strand of each coding sequence in each library. The PPB measures the normalized rNMP-embedment frequency at each base (see **Methods**). Each row represents a CDS, which is sorted by descending gene size. Each column represents an rNMP library. There are three control libraries with artificial rNMP embedment generated *in silico* on the right (see **Methods**). **(B)** Linear regression plot showing a positive relationship between the rNMP-enrichment factor and the gene size on the non-template strand. Each data point represents the rNMP-enrichment factor on the non-template strand of a particular gene in an rNMP library of a given cell type. The rNMP-enrichment factors are calculated as described in **Methods**. Spearman’s r and p-value are marked in the plot. Cell types with Spearman r > 0.5 have a strong positive correlation between the rNMP-enrichment factor and the gene size. The 95% confidential interval is marked as the blue shadow region around the line.

Besides the 13 CDSs, there are 2 rRNA and 22 tRNA genes in the hmtDNA. The longer rDNA (MT-RNR2) has a higher rNMP-embedment frequency than the shorter one (MT-RNR1) in all libraries on the non-template strand. The tRNA genes have similar lengths ranging from 58 to 74 nt. Given the short length of tRNA genes, few rNMPs are incorporated, which leads to high variation in the frequency and non-significant association between tRNA gene length and rNMP embedment frequency (**Additional file 3: Fig. S5D**).

### The composition of incorporated rNMPs in hmtDNA of different cell types

The rNMP-embedment analysis of the 32 hmtDNA libraries reveals that rUMPs are the least abundant rNMPs in all cell types despite the strand preference and RNase H2 genotype (**Fig. 5A**). However, the preferred rNMPs are different among the cell types. In CD4^+^T, liver-tissue, and HCT116 libraries, rAMPs are preferred. In hESC-H9, WB-GTP control, WB-GTP PTSD, and HEK293T RNH2A WT libraries, rCMPs are preferred. In the HEK293T RNH2A-KO libraries, both rCMPs and rGMPs are preferred (**Fig. 5A and Additional file 3: Fig. S7A**). We located the high-frequency rNMP sites (Top 1% sites having the most abundant embedded rNMPs, see **Methods**) in mtDNA and analyzed their sequences. These high-frequency rNMP sites also show similar preferred rNMPs as in the whole hmtDNA (**Additional file 3: Fig. S8**, 0 position is the rNMP position). CD4^+^T, liver tissue, and HCT116 cells show preferences for rAMPs in most libraries, whereas hESC-H9 libraries have a strong preference for rCMPs. HEK293 RNH2A WT and KO cells show preferences for rCMPs and rGMPs.

**Figure 5.**
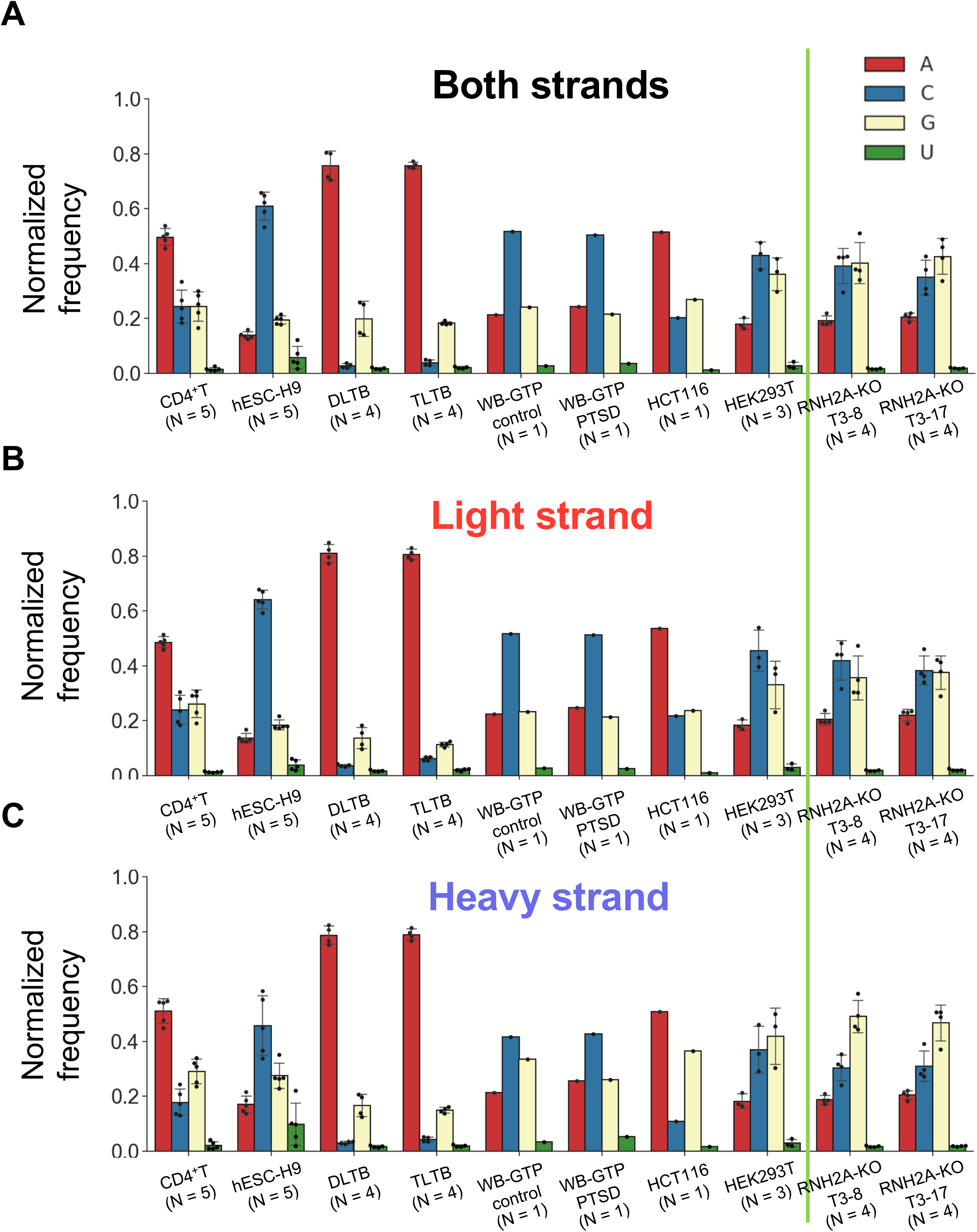
The rNMP composition on both strands of hmtDNA.

Bar plot showing the normalized frequency of the four rNMP types on **(A)** both strands, **(B)** light strand, and **(C)** heavy strand of hmtDNA. The normalized frequency is calculated as described in the **Methods** section. Types of rNMPs and their corresponding colors are listed in the color legend on the right top. The vertical, green line separates the cell types with RNH2A-WT and with RNH2A-KO genotype.

Our analysis reveals a slight difference in rNMP composition between the light and heavy strands of hmtDNA. The heavy strands have more rGMPs incorporated, especially in HEK293T WT and RNH2A-KO libraries (**Fig. 5B, C and Additional file 3: Fig. S7B, C**). This difference does not exist between forward and reverse strands of yeast mtDNA (**Additional file 3: Fig. S9**). Moreover, the rNMP compositions do not match the rNTP abundancy relative to that of the dNTPs in the cells (**Additional file 3: Fig. S10**), which suggests that the rNTP and dNTP pools are not the only determinants for rNMP-embedment composition and patterns, and rNMP-embedment may be related to other cellular functions.

### Genomic context of the embedded rNMPs

Distinct dinucleotide patterns consisting of the incorporated rNMP and its direct upstream dNMP neighbor are revealed in this study. Specifically, TrA, CrC, and ArG are preferred in all cell types while CrA and CrU are preferred in all cell types except liver-tissue cells (**Fig. 6A**, P < 0.05 in **Additional file 6**). Compared to other cell types, liver-tissue cells have more dGMPs located upstream of all four types of rNTPs (**Fig. 6A**).

**Figure 6.**
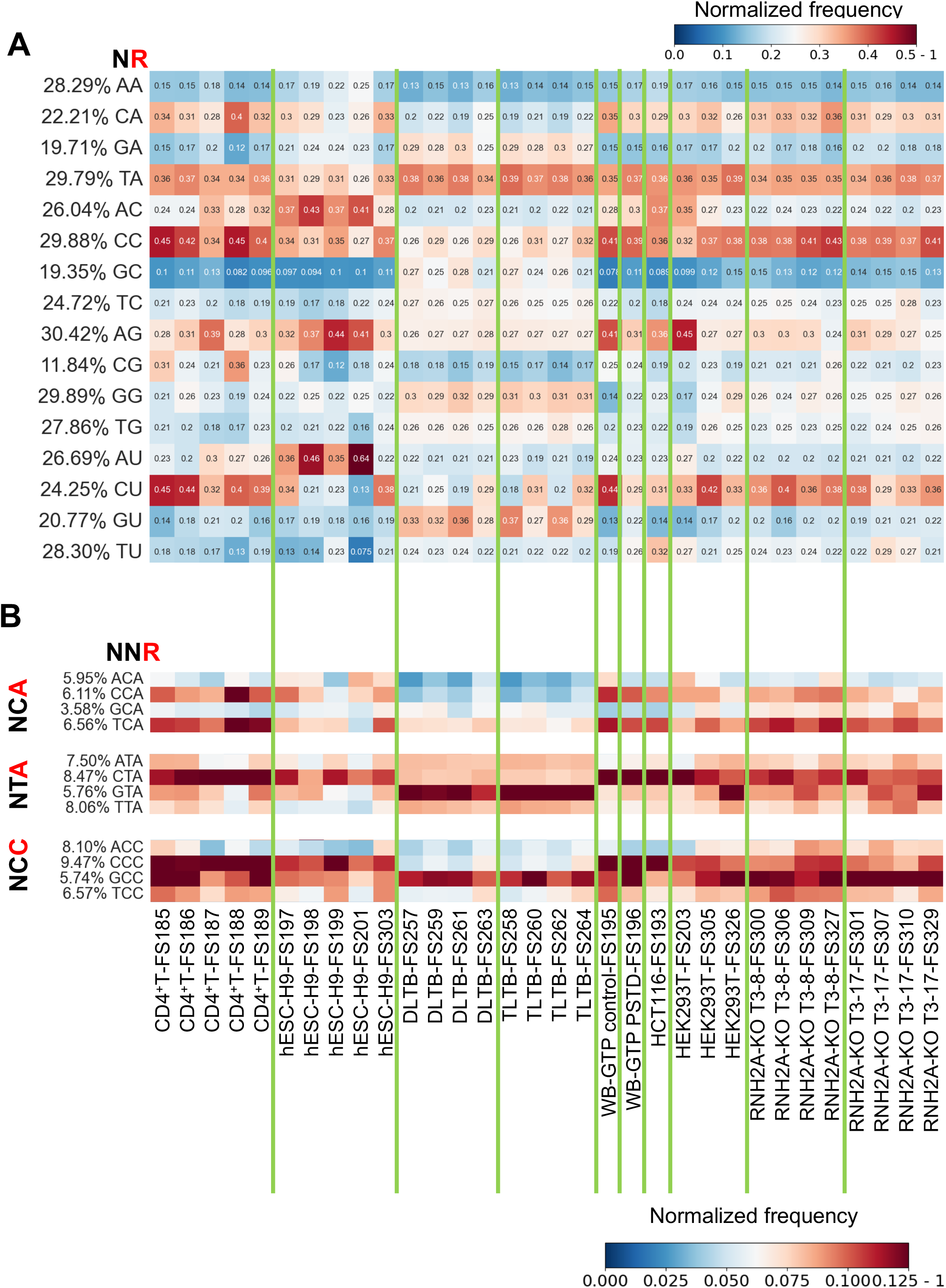
Preferred rNMP embedment patterns in hmtDNA. (**A**) Heatmap analyses with the normalized frequency of dinucleotides composed of the incorporated rNMP (R: rA, rC, rG, or rU) and its upstream dNMP neighbor (N: dA, dC, dG, or dT) (NR) in the hmtDNA. The rNMP position in the dinucleotide is shown in red at the top left of the heatmap. Each column of the heatmap shows the results of an rNMP library. The vertical, green lines separate different cell types. The color scale is shown in the top right corner of the figure. **(B)** Heatmap analyses with the normalized frequency of selected trinucleotides composed of the incorporated rNMP (R: rA, rC, rG, or rU) and its two upstream dNMP neighbors (N: dA, dC, dG, or dT) (NNR) in the hmtDNA. The color scale is shown in the bottom right corner of the figure.

Similar to what was observed in yeast mitochondrial and nuclear DNA (21), there are fewer dinucleotide patterns formed by rNMPs and their downstream dNMP (RN) due to the downstream dNMP being incorporated following the incorporation of the rNMP. The only two RN patterns we identified were rAG and rUA in liver-tissue samples. There is no strong RN pattern in other cell types (**Additional file 3: Fig. S11**).

Our analysis reveals that there is strand bias for particular dinucleotide patterns (NR). We found that CrU is stronger on the light strand in the HEK293T RNH2A-KO libraries while CrC is stronger on the heavy strand in the liver-tissue and HEK293T RNH2A-KO libraries (**Additional file 3: Fig. S12,** P < 0.05 in **Additional file 7**). In contrast, the forward and reverse strands of mtDNA have quite similar dinucleotide patterns in yeast. CrA, ArC, ArG, and CrU are consistently preferred in both strands of yeast RNase H2 WT and *rnh201*-null libraries ((21) and **Additional file 3: Fig. S13**).

Furthermore, we identified preferred trinucleotide patterns in hmtDNA. GTrA and GCrC are strong in all cell types but preferred in liver-tissue libraries. CCrA, TCrA, CTrA, and CCrC are preferred in all libraries except liver-tissue libraries. (**Fig. 6B and Additional file 3: Fig. S14A**, P < 0.05 in **Additional file 5**). The light and heavy strands have many different preferred trinucleotide patterns. For example, there are stronger CCrA and CCrG patterns on the light strand, and more AArG and ATrG on the heavy strand. (**Additional file 3: Fig. S14**). In the yeast libraries, there are biased trinucleotide frequencies, including stronger GTrC on the forward strand and stronger CCrA on the reverse strand (**Additional file 3: Fig. S15**).

## Discussion

Through the analysis of 32 libraries of rNMP sites embedded in mtDNA of 6 different human cell types, the results of this study support non-random rNTP presence in human mtDNA. We found an rNMP-embedment bias on the light strand of hmtDNA of most cell types examined except for the liver-tissue samples (**Fig. 1A and Additional file 3: Fig. S1**). We identified common zones with enriched rNMP presence in hmtDNA, which we term REZs (**Fig. 1C and Additional file 4**). Among all of our mtDNA samples, we found eight common REZs on the light strand and five common REZs on the heavy strands (**Fig. 1C**). These common REZs are present in the hmtDNA of all six cell types studied independently from the RNase H2 genotype. The REZs are not linked to the method of DNA fragmentation used. Albeit the different fragmentation methods lead to variant rNMP capture efficiency, all of our libraries show rNMP enrichment in the same REZs. REZs also exist in yeast mtDNA. By analyzing the rNMP distribution in mtDNA of published yeast rNMP libraries constructed with two distinct rNMP-mapping techniques (ribose-seq (22) and emRiboSeq (25)), we found that all of these yeast libraries contain the same REZs in yeast mtDNA (**Additional file 3: Fig. S1B**). These results corroborate the nonrandom distribution of rNMPs in mtDNA and highlight the presence of DNA regions displaying rNMP accumulation. The REZs may be closely related to mtDNA structure and function, and need further investigation.

In this study, we identified three common REZs in the D-loop of the control region in all cell types (**Fig. 3A, B and Additional file 3: Fig. S3**). Interestingly, in the same zones, mtDNA polymerase Pol γ and mtDNA helicase TWINKLE show lower binding activity compared to other regions in the D-loop (**Additional file 3: Fig. S4**). Considering that the incorporated rNMPs can alter the DNA structure and affect DNA-protein interaction (7, 23), the low binding activity of Pol γ and TWINKLE may be induced by the enrichment of rNMPs. Since Pol γ and TWINKLE are essential for hmtDNA replication, these REZs may be one possible cause or consequence of frequent mtDNA replication failure (38). In hmtDNA, most of the initiation events are prematurely terminated, generating the 7S DNA product (38). Frequent rNMP embedment in the control region may be associated with this phenomenon. Compared to the other cell types analyzed, liver-tissue cells have a stronger heavy-strand REZ in the control region, which may be related to the abundance of the 7S DNA (**Fig. 3A, B and Additional file 3: Fig. S3**).

Beyond a connection between rNMP presence in DNA and replication in hmtDNA, we found that rNMP embedment is associated with genes and transcription in hmtDNA. The two strands of hmtDNA have clear distinct functions in mtDNA transcriptions (35). Thus, the consistent strand preference of the incorporated rNMPs on the light strand in most cell types and on the heavy strand of liver-tissue cells may play a role in hmtDNA transcription (**Fig. 1**). We found a significant positive correlation between CDS length and rNMP-embedment frequency on the non-template strand (**Fig. 4**). We suppose that this phenomenon may be related to the CDS sequence or mtDNA structure. During transcription, mRNA may displace the non-template strand and form R-loops, which may lead to the incorporation of parts of the RNA strands into the DNA of the non-template strand. Longer CDSs may have more R-loops generated during transcription (48). It would be interesting to determine whether these sites of embedded rNMPs in the long CDSs are due to rNMP tracts rather than single rNMPs. Moreover, embedded rNMPs may contribute to premature transcription termination in the OriH region. The three conserved sequence blocks (CSB1: 213-235, CSB2: 299-315, CSB3: 346-363) located in the human OriH region (nucleotides 110-441) are highly conservative sequences located in the control region of mtDNA in many vertebrate species, and frequent premature termination of light-strand transcripts have been observed in the CSBs (49, 50). We found strong light-strand rNMP-embedment preference in the OriH region of all cell types including liver-tissue cells, in which 80% of rNTPs are incorporated on the heavy strand (**Fig. 3C, D**). As shown in a recent study, the embedded rNMPs can impair RNA polymerase activity and lead to transcription termination (39). The high rNMP-embedment rate on the light strand may affect the POLRMT binding activity and contribute to premature transcription termination.

In hmtDNA, rUMP is the least abundant rNMP detected in all cell types studied after normalization to the nucleotide content of the hmtDNA genome and independently from the RNase H2 genotype of the cells. This is in common with the rNMP composition in mtDNA of other eukaryotic cells, such as budding (*S. cerevisiae* and *S. paradoxus*) and fission (*S. pombe*) yeast (21), and the unicellular green alga (*C. reinhardtii*) (34). As observed in yeast and algae, the rUTP/dTTP ratio calculated from the concentrations of all rNTPs and dNTPs in the pool of the human cells analyzed is the smallest, while the rATP/dATP and rGTP/dGTP are the largest ratios **(Fig. 5 and Additional file 3: Fig. S10**). However, in contrast to what was observed in *C. reinhardtii*, in which rAMP was by far the most abundant rNMP in the mtDNA and chlDNA of the alga in line with a large rATP/dATP ratio, the rNTP/dNTP ratios of the bases A, C, and especially G, do not match the rNMP composition in the hmtDNA (34). This inconsistency suggests that human mtDNA Pol γ may have a preference for rNTP incorporation in hmtDNA. The relatively lower rGTP incorporation rate compared to its abundance may in part be due to rGTP’s frequent interaction with proteins in human cells (51).

Previous studies in yeast and algae showed that the rNTPs are preferentially incorporated after a particular type of dNMP in DNA (dA, dC, dG, and dT) (21, 34, 52). In hmtDNA, we identified several preferred dinucleotide and trinucleotide patterns. We found TrA and CrC to be the most conserved patterns in hmtDNA among all cell types tested (**Fig. 6A**). These dinucleotides are preceded by dG (GTrA and GCrC) in the liver-tissue samples, while they are preceded by dC (CTrA and CCrC) in all other cell types (**Fig. 6B**). This dG and dC bias are in line with the more abundant rNMP presence on the heavy strand (G-rich) for the liver-tissue samples as opposed to the light strand (C-rich) for the other cell types (**Fig. 1A**). Among the strongly conserved CrA, ArC, ArG, and CrU pattern of yeast mtDNA, only CrU and CrA are found in hmtDNA and not in all cell types. Interestingly, the rNMP patterns found in hmtDNA are more similar to those of the fission yeast than those of budding yeast, with a strong CrC pattern (21). Since RER does not work to remove rNMPs in hmtDNA, the rNMP-embedment pattern in hmtDNA mainly reflects the rNTP-incorporation pattern of human Pol γ. The different rNMP embedment patterns may thus reflect a distinct mtDNA polymerase structure in these different species.

## Conclusions

This study reveals rNMP-embedment characteristics in hmtDNA across six different human cell types including cancer samples and compares them with those found in yeast mtDNA. Our analyses support the existence of rNMP-enriched zones (REZs) in hmtDNA, which are surprisingly conserved among the different cell types and potentially function-related. We found three REZs in the replication control region of hmtDNA, which may affect the binding activity of DNA Pol γ and TWINKLE, whereas the strong light-strand preference in the OriH region in all cell types is potentially related to transcription termination. All of our findings indicate that the rNMP presence in hmtDNA is not random. There is marked strand bias of rNMP distribution, which is dependent on the cell types analyzed. While rUMP is always the least frequent rNMP incorporated in hmtDNA, different cell types have different preferred rNMPs. There are rNMP hotspots and patterns, which are either common or specific to the cell types tested. Moreover, there is a higher rNMP-embedment frequency per nucleotide in longer coding sequences in mtDNA in all cell types. Altogether, our findings are laying the groundwork to understand the biological and medical relevance of RNA intrusions in the DNA of normal and diseased cells.

## Methods

### Human cell line preparation

#### Human primary activated CD4^+^ T cells

CD4^+^T cells were isolated from human buffy coats of 5 health donors (NY Blood Center) as previously described (53), and the isolated CD4^+^T cells were pooled and activated by phytohemagglutinin (5ng/ml) and IL-2 (5ng/ml) for 3 days. The total cellular DNAs of the activated CD4+ T cells were isolated using a DNA extraction kit (Promega Wizard) for analysis.

#### hESC-H9

Human Embryonic Stem Cells (hESC) line H9, passage 22, were purchased from WiCell Research Institute, University of Wisconsin (line WA09, lot # WB66595). hESC was maintained and expanded in feeder-free culture conditions with mTeSR basal medium (STEMCELL Technologies Inc., Vancouver, Canada, cat # 85850) on 6-well plates culture-treated coated with Corning Matrigel hESC-qualified Matrix, LDEV free (Corning Inc., Corning, NY, USA, catalog # 35427) in a humidified chamber in a 5% CO_2_-air mixture at 37°C. The culture medium was changed daily, and regions of differentiation were removed by aspiration. Cells were passaged as small aggregates every four to five days at ratios of 1:3 to 1:6 using a gentle dissociation reagent (Corning Inc., cat # 3010). Cells were harvested at passage 28 as a single-cell suspension for downstream applications.

#### DLTB and TLTB

Paired HCC Tumor Liver Tissue Biopsy (TLTB) and adjacent non-tumor Distal Liver Tissue Biopsy (DLTB) from patients undergoing HCC resection were obtained from the Department of Medicine, General Surgery and Transplantation of the University of Udine, Udine, Italy. None of the patients had received any local or systemic anticancer treatments before the surgery. Both tumor and non-tumor tissues were histologically confirmed. Mitochondria were isolated from surgical biopsies as reported by Bazzani et al (54). Then, mtDNA was extracted and purified using Qiagen genomic-tip 20/G and following the manufacturer’s indications. DLTB and TLTB samples of 17 patients affected by HCC and characterized by the cirrhotic liver due to alcohol abuse or HCV and HBV infection were pooled together to obtain a sufficient amount of mtDNA to construct the ribose-seq library. The individual DLTB-8 and TLTB-8 samples were obtained from a 68 years old Caucasian male, who was diagnosed with a large HCC located in the right liver lobe during a routine ultrasound. A computed tomography scan confirmed the diagnosis. Differently from the pooled sample, the patient had no history of alcohol intake and resulted negative for HCV and HBV. Liver function was normal without signs and symptoms of portal hypertension (Child-Pugh score A-6 and MELD 7). His preoperative alpha-fetoprotein (AFP) was 4.8 ng/ml. He underwent an eventful open right hepatectomy in 2015; during surgery, the liver resulted macroscopically normal without cirrhosis. Pathological examination revealed a well-differentiated HCC (G1) with a maximum diameter of 9.5 cm, microvascular invasion was absent resulting in a pathological staging pT1, N0, M0. The patient was followed-up at the Institution yearly since the last follow-up in April 2022 when he resulted negative for HCC recurrence, with normal liver function and an AFP of 4.6 ng/ml.

#### WB-GTP

Whole Blood Grady Trauma Project (WB-GTP) was part of a larger investigation of genetic and environmental factors in a predominantly African American (AA) urban population of low socioeconomic status with the purpose of determining how those factors may modulate the response to stressful life events. Research participants were approached in the waiting rooms of primary care of a large, public hospital (Grady Memorial Hospital in Atlanta, Georgia) while either waiting for their medical appointments or while waiting with others who were scheduled for medical appointments. Screening interviews, including the participants’ demographic information (e.g., self-identified race, sex, and age) and psychiatric history were completed on-site, including the Clinician Administered PTSD Scale (CAPS (59)). Subjects were scored as having PTSD if they met DSM-IV PTSD criteria from the CAPS interview. Exclusion criteria included mental retardation or active psychosis. Further details regarding the Grady Trauma Project (GTP) dataset can be found in Gillespie et al (60). Written and verbal informed consent was obtained for all subjects. DNA from the participant’s blood was extracted using either the E.Z.N.A. Mag-Bind Blood DNA Kit (Omega Bio-Tek, Inc., Norcross, GA) or ArchivePure DNA Blood Kit (5 Prime, Inc, Gaithersburg, MD) following protocol instructions. Matching samples with the highest yield were selected for the analyses. All procedures in this study were approved by the Institutional Review Boards of Emory University School of Medicine and Grady Memorial Hospital.

#### HCT116

HCT116 cells were provided by the Pursell group at Tulane University. The cells were grown in Dulbecco’s modification of Eagle’s medium (DMEM) containing non-essential amino acids (Corning) with 10% fetal bovine serum (Sigma-Aldrich) and 1x penicillin-streptomycin (Gibco). Cells were grown at 37 °C in a 5% CO_2_-humified incubator. Cells were removed from the plate by 3-5 min incubation in media containing trypsin. Cells were spun down and the supernatant was removed. Cells were lysed by the addition of TNES lysis buffer containing Proteinase K followed by phenol: chloroform extraction. Genomic DNA was ethanol precipitated, air dried, and concentration quantified using a Nanodrop and Qubit fluorometer (Thermo Fisher Scientific).

#### HEK-293T

Human embryonic kidney T (HEK-293T) RNASEH2A wild-type and KO (RNH2A-KO) cells were provided by the Pursell group at Tulane University. The cells were grown in Dulbecco’s modification of Eagle’s medium (DMEM) containing 4.5 g/L glucose, L-glutamine, and sodium pyruvate (Corning) with 10% fetal bovine serum (Sigma-Aldrich) and 1x penicillin-streptomycin (Gibco). Cells were grown at 37 °C in a 5% CO_2_-humified incubator. For the construction of the RNH2A KO clones, RNASEH2A (Chr19, exon 2) gRNA (5’-TAACAGATGGCGTAGACCAT-3’) was cloned into GeneArtTM CRISPR Nuclease Vector with OFP reporter (Invitrogen) following manufacturer’s protocol. HEK-293T cells were transfected with 6 mL of Lipofectamine 2000 (Invitrogen) and 2.5 mg of gRNA vector DNA when the cells were 60-70% confluent. After 48 hours, OFP-positive cells were FACs sorted and serially diluted into 96-well plates at ∼1.5 cells per well and incubated for 10-14 days. HEK-293T RNH2A-KO T3-8 and T3-17 have three distinct frameshift mutations consistent with all three alleles being modified in hypotriploid 293T cells, respectively. The RNH2A-KO T3-8 has an insertion of G at position -1; deletion of five bases at position +2 to +6; complex alteration with deletion of three bases from position -1 to +2, deletion of 2 two bases from position +5 to +6, and CC>TT at position +8 and +9. The RNH2A-KO T3-17 has the deletion of 14 bases from position + 7 to -7; the deletion of six bases from position -5 to 1; and the deletion of 25 bases from position -3 to +22. All positions are indicated with respect to the Cas9 cleavage site on the reference strand.

### Ribose-seq library preparation

The libraries in this study were prepared using the latest ribose-seq method that has been upgraded from the previous method (21, 22, 55). To construct libraries with human cell lines, we improved the ribose-seq protocol by i) introducing NEBNext® dsDNA Fragmentase® (New England Biolabs) to fully fragment the human genomic DNA samples; ii) optimizing the removal of linear ssDNA process to retain more rNMPs; iii) applying a new size selection method using HighPrep™ PCR Clean-up System (MagBio Genomics) to increase the yield of captured rNMPs.

All the commercial enzymes applied in the ribose-seq protocol were used according to the manufacturer’s recommended instructions. Genomic DNA samples were fragmented using NEBNext® dsDNA Fragmentase® or a combination of restriction enzymes to generate small DNA fragments. Various sets of restriction enzymes were applied to fragment human genomic DNA samples, as indicated in **Additional file 1**. The used combinations were i) RE1: [HpyCH4V] + [Hpy116II, Eco53KI, RsaI, and StuI]; ii) RE2: [AleI, AluI, and PvuII] + [DraI, HaeIII, and SspI]; iii) RE3: [CviKl-1] + [Mly, MscI, and MslI].

Following fragmentation with restriction endonucleases, the fragmented DNA was purified by QIAquick PCR Purification Kit (Qiagen). The fragments were tailed with dATP (New England Biolabs) by using Klenow Fragment (3′→5′ exo-) (New England Biolabs) for 30 min at 37 °C and purified by using a QIAquick PCR Purification Kit. In case of fragmentation with NEBNext® dsDNA Fragmentase®, NEBNext End Repair Module (New England Biolabs) was performed before dA-tailing to convert fragmented DNA to blunt-ended DNA having 5′ phosphates and 3′-hydroxyls.

Following dA-tailing and purification, a partially double-stranded adaptor (Adaptor. L1 - Adaptor.L8 with Adaptor.S (21)) was annealed with the DNA fragments by T4 DNA ligase (New England Biolabs) incubating overnight at 16 °C. Following overnight ligation, the products were purified using HighPrep™ PCR Clean-up System.

The adaptor-annealed fragments were then treated with 0.3 M NaOH (Working concentration) for 2 h at 55 °C to denature the DNA strands and cleave the 3’ site of embedded rNMP sites resulting in 2′,3′-cyclic phosphate, and 2′-phosphate termini.

After the treatment with Alkali, neutralization using 2 M HCl and purification using HighPrep™ RNA Elite Clean-up System (MagBio Genomics) was performed. Further purification steps were performed using HighPrep™ RNA Elite Clean-up System, except for the size selection step.

The single-stranded DNA (ssDNA) fragments were then incubated with 1 μM *Arabidopsis thaliana* tRNA ligase (AtRNL), 50 mM Tris-HCl pH 7.5, 40 mM NaCl, 5 mM MgCl_2_, 1 mM DTT, and 300 μM ATP for 1 h at 30 °C to ligate the 5’-phosphate and 3’-OH of ssDNA fragments containing an rNMP, followed by bead purification.

The ssDNA fragments were treated with T5 Exonuclease (New England Biolabs) for 30 min at 37 °C to degrade the unligated linear ssDNA fragments. After purification, the circular fragments were incubated with 1 μM 2′-phosphotransferase (Tpt1), 20 mM Tris-HCl pH 7.5, 5 mM MgCl_2_, 0.1 mM DTT, 0.4% Triton X-100, and 10 mM NAD^+^ for 1 h at 30 °C to remove the 2′-phosphate at the ligation junction.

Following purification, the circular ssDNA fragments were amplified with two steps of PCR and become a library. Both PCR began with an initial denaturation at 98 °C for 30 s. Then denaturation at 98 °C for 10 s, primer annealing at 65 °C for 30 s, and DNA extension at 72 °C for 30 s were done. These PCR steps were performed for 6 cycles in the first PCR round, and for 11 cycles in the second PCR round. Lastly, there is a final extension reaction at 72 °C for 2 min for both PCRs.

A first round of PCR was performed to extend the sequences of the Illumina adapter for TruSeq CD Index primers. The primers (PCR.1 and PCR.2) used for the first round were the same for all libraries. A second round of PCR was performed to insert specific indexes i7 and i5 into each library. Both PCR rounds were performed with Q5-High Fidelity polymerase (New England Biolabs) for 6 and 11 cycles, respectively.

Following the PCR cycles, DNA fragments between 250 and 700 bp were purified using the HighPrep™ PCR Clean-up System. The resulting ribose-seq libraries were mixed at equimolar concentrations and normalized to 4 nM. The libraries were sequenced on an Illumina Next 500 or Hiseq X Ten in the Molecular Evolution Core Facility at the Georgia Institute of Technology or Admera Health.

### Locate rNMPs in DNA

Raw sequenced reads are trimmed using Trim_galore software with the command “trim_galore - a AGTTGCGACACGGATCTATCA -q 15 --length 62 input.fastq -o output.fastq”. Using this command, the remaining adapter sequences and low-quality bases with sequencing quality scores lower than 15 are trimmed. Afterward, the Ribose-Map bioinformatics toolkit is used to map the incorporated rNMPs to the human genome GRCh38 mitochondrial DNA sequence (chrM) on a single-nucleotide level (30, 36). Since Ribose-Map processes hmtDNA as a linear DNA, we need to recover the rNMP embedment around the zero position. Specifically, we used Ribose-Map to align the rNMPs to generate control-region rNMPs. Then, control-region rNMPs and linear hmtDNA rNMPs are merged to form the actual rNMPs in circular hmtDNA. Afterward, the background noise of RE-recognition sites is removed using the same protocol in (21). Some rNMPs mismatch the nucleotide at their alignment locations in the reference genome. These rNMPs are likely to be induced by sequencing error and are also removed from all libraries using customized python3 scripts available under GNU GPL V3.0 License on GitHub (https://github.com/xph9876/rNMP_match_analysis).

### Calculate the normalized frequencies for rNMP embedment

We use the rNMP-embedment probability per base (PPB) and enrichment factor to measure the rNMP-embedment frequency in a particular region. The PPB represents the rNMP-embedment probability at each nucleotide, which is comparable among the different genomic regions and different rNMP libraries. For genomic region *G* in library *L*, the PPB can be calculated using the following formula.

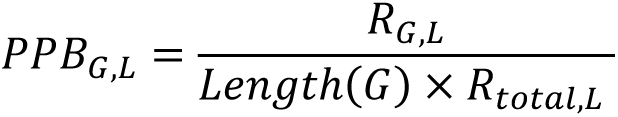

*R_G,L_* denotes the rNMP count in the genomic region *G* in library *L*, and *R_total,L_* denotes the total rNMP count in mtDNA in library *L*. The raw PPB value is used to measure the rNMP embedment frequency in coding sequence comparison. The moving average of PPB (window size = 51 nt) is used to measure the rNMP embedment frequency in the control region. The enrichment factor is similar to PPB but measures the rNMP embedment frequency in a region, which can be calculated as

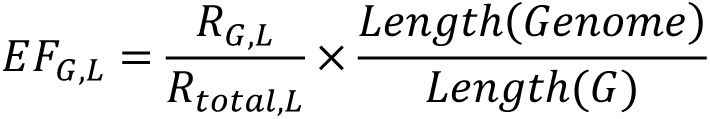

The normalized frequency of the rNMP composition and rNMP patterns is calculated as described in Balachander et al (21). Briefly, the count of rNMPs, dinucleotides, and trinucleotides are divided by the corresponding background dNMP count in the reference genome to get the frequency. For composition, the frequencies are further divided by the sum of frequencies to get the normalized frequencies. For dinucleotide and trinucleotide patterns, the frequencies are divided by the sum of frequencies that share the same rNMP to get the normalized frequencies. The normalized frequency is calculated using the RibosePreferenceAnalysis package (33).

### Determine the rNMP hotspots and high-frequency rNMP locations in mtDNA

To identify the rNMP hotspots for a specific cell type, or all cell types combined, we searched the reference genome to find the single-nucleotide locations with incorporated rNMPs in all libraries using a customized R script. To identify high-frequency rNMP locations, which are the locations with the highest rNMP incorporated in a single base of hmtDNA, we calculated the frequency of rNMP counts at single-base locations for every library at each location on the chromosome with the presence of at least 1 rNMP. We then filtered out the top 1% locations with the most abundant embedded rNMPs and extracted +/− 3 bp genomic sequences around the rNMP locations to generate a consensus sequence using the ggseqlogo R package.

### Detect rNMP enriched zones (REZs) in hmtDNA

Both strands of hmtDNA were divided into 200-nt bins (N = 166). For each rNMP library, the rNMP enrichment factors (EF) were calculated in each bin. All bins with EF > 1 represent enriched zones in the library. The common REZs are the bins that are enriched in all cell categories and at least in 80% of the libraries.

### Background coverage calculation for different fragmentation methods

Paired-end sequencing was performed on fragmented CD4^+^T cell DNA using restriction enzymes and/or dsDNA fragmentase. Paired-end sequencing reads were aligned to circular hmtDNA with a maximum insert size of 1000 nt. Afterward, coverage at each nucleotide in hmtDNA was calculated by piling up the aligned reads. The enrichment factor of coverage inside each 200-nt bin was calculated as background coverage.

### Generate random rNMP library for control

Three rNMP libraries of 10,000, 100,000, and 1,000,000 rNMPs randomly incorporated in hmtDNA were used as the control of the CDS study.

### Mass spec analysis of nucleotides

dNTP and rNTP concentration (pmol/million cells) for activated CD4^+^T cells were collected from a previous study having 4 replicates, and the precalculated average was used to represent dNTP/rNTP ratios (56). The nucleotide samples were extracted based on the established protocol (57) with some modifications. To prepare the nucleotide samples, 2 x 10^6^ cells of HEK293T or hESC-h9 were counted and centrifugated to obtain a cell pellet. The pellet was washed with PBS and then vortexing was performed for 2 min with 200 µL of cold 65 % methanol for cell lysis. The cell mixture was incubated at 95 °C for 3 min and then incubated on ice for 1 min to complete the cell lysis. By centrifugation at 14,000 rpm for 3 min, the supernatant containing nucleotides was isolated. To quantify the intracellular dNTPs and rNTPs, an ion pair chromatography-tandem mass spectrometry method (57) was applied, with modifications. Chromatographic separation and detection were performed on a Vanquish Flex system (Thermo Fisher Scientific) coupled with a TSQ Quantiva triple quadrupole mass spectrometer (Thermo Fisher Scientific). Analytes were separated using a Kinetex EVO-C18 column (100 × 2.1 mm, 2.6 µm) (Phenomenex) at a flow rate of 250 µL/min. Pmol/million cells were calculated for rNTPs and dNTPs for all four replicates to calculate the ratios.

### Human genes in the study

The human gene annotation GTF file was downloaded from the human reference genome GRCh38 in Ensembl genome browser 108. The mtDNA genes were selected from the GTF file. CDS and non-coding genes were also indicated in the same file. The processed gene locations are available on GitHub (https://github.com/xph9876/rNMP_hmt_analysis).

### Compare light and heavy strand sequencing bias using DNA sequencing

Whole genome sequencing data of genomic DNA samples extracted for ribose-seq were also obtained using the Illumina NextSeq 500 System with Mid-output (PE150) in the Molecular Evolution Core Facility at the Georgia Institute of Technology. The FASTQ reads were trimmed to remove any residual Illumina adapters and filtered to keep reads above the quality threshold of 15 and length 50bp. These trimmed reads were then aligned using Bowtie 2 (58) with the hg38 reference genome. The reads aligned to the light and heavy strands of mtDNA were calculated to check any strand-biased sequencing. The coverage-mean depth was calculated per base pair using SAMtools and was greater than 30x for all libraries.

### Statistical test

To test if a dinucleotide or trinucleotide pattern was preferred in human or yeast mtDNA, a one-tailed Mann-Whitney *U* test was used to compare the normalized frequency for each dinucleotide or trinucleotide pattern and the expectation value (0.25 for dinucleotide and 0.0625 for trinucleotide). To test if one dinucleotide or trinucleotide pattern was preferred on the light or the heavy strand, a two-tailed Mann-Whitney *U* test was performed. For both statistical tests, the significance level of 0.05 was used.

### Visualizations

Alignment of ribose-seq data and finding the coordinates was done using Ribose-Map (30, 59). The rNMP composition and dinucleotide heatmaps were generated using RibosePreferenceAnalysis (33). All other figures were generated using Python3 scripts, which are available on GitHub (https://github.com/xph9876/rNMP_hmt_analysis). Detection of common and high-frequency rNMP locations with ggseqlogo plot visualizations was conducted using R scripts available on GitHub (https://github.com/DKundnani/rNMP_hotspots).

## Declarations

### Ethics approval and consent to participate

The DLTB and TLTB study was approved by the Unique Regional Ethic Committee, on August 24, 2019, Protocol number 18659, and informed consent was obtained from each patient.

Studies on the hESC-H9 stem cells are approved by the Institutional review board (IRB), protocol number H18243.

Written and verbal informed consent was obtained for all participants, and all procedures in this study were approved by the institutional review boards of Emory University School of Medicine and Grady Memorial Hospital, Atlanta, Georgia.

### Consent for publication

Not applicable.

### Availability of data and materials

The DNA-seq and ribose-seq datasets for all hmtDNA libraries are available in NCBI’s BioProject via accession number “PRJNA941970”. The ribose-seq dataset for budding yeast *S. cerevisiae* is available in NCBI’s BioProject via accession number “PRJNA613920”. The emRiboSeq datasets analyzed are available in NCBI’s Gene Expression Omnibus via accession number “GSE64521”. Codes generated and used for analysis are available under MIT License on GitHub (https://github.com/xph9876/rNMP_hmt_analysis). The scripts used to remove rNMPs at restriction enzyme locations are available under the GNU GPL V3.0 license on GitHub (https://github.com/xph9876/ArtificialRiboseDetection). The scripts used for rNMP mismatch removal are available under GNU GPL V3.0 License on GitHub (https://github.com/xph9876/rNMP_match_analysis). The scripts used for locating rNMP hotspots and high-frequency rNMP embedment positions are available under the MIT License on GitHub (https://github.com/DKundnani/rNMP_hotspots).

## Authors’ contributions

F.S. together with P.X. and T.Y. conceived the project and designed experiments with the help of G.T. F.S. and P.X. wrote the manuscript with help from T.Y., D.L.K., and M.S. T.Y. performed most of the experiments with help from S.M., Y.J., G.N., and S.B. P.X. performed most of the bioinformatics analyses with the help of D.L.K., M.S., and A.L.G. S.M. performed the experiment with the hESC-H9 samples. V.B., U.B., and C.V. provided DLTB and TLTB samples. V.S.P. and Z.F.P. constructed the RNase H2A KO cell lines. A.L. provided the WB-GTP samples. B.K. provided the CD4^+^T samples. S.T., R.F.S., and B.K. performed the measurements of dNTPs and rNTPs. All authors commented on and approved the manuscript.

## Supporting information

All additional files

## Acknowledgments

We thank S. Biliya, N. Djeddar, and A. Bryksin from the Molecular Evolution Core for advice and support with high-throughput sequencing, and the Partnership for an Advanced Computing Environment (PACE) at the Georgia Institute of Technology for their research cyberinfrastructure resources and services. We thank A. Smith at Emory University for providing the WB-GTP samples. We are grateful to the staff and participants from the Grady Trauma Project for their time and effort in supporting this research. We acknowledge T. Channagiri, F. M. Figueroa, Y. Lee, and S. Randhawa for critically reading the manuscript; and all members of the Storici laboratory for assistance and feedback on this study.

