## Supplementary material for "Enriched zones of embedded ribonucleotides are associated with DNA replication and coding sequences in the human mitochondrial genome": All additional files: additional file 3 Supplementary table and figures 20230405.pdf

**Additional file 1. rNMPs found in ribose-seq libraries of hmtDNA.**

List of mitochondrial ribose-seq libraries constructed in this study with corresponding data information and an indication of cell type, genotype, library name, fragmentation method, number of rNMPs, % rA, rC rG, and rU with mean and standard deviation for each cell type, barcode, and number of cycles in PCR 1 and PCR 2 of the ribose-seq protocol.

**Additional file 2 Counts and percentages of reads aligned to light and heavy strands of hmtDNA.**

The table shows the number of reads aligned to hmtDNA for DNA sequencing data from the genomic DNA of cell lines used for ribose-seq in this study. Percentages for both strands are calculated from the read count. The last column shows the mean depth of coverage per base suggesting enough coverage of at least 40x coverage to determine the strand bias.

**Additional file 3 Supplementary table and figures.**

Supplementary Table 1 and Supplementary Figures 1-15 of this study.

**Additional file 4 Common rNMP-enriched zones in hmtDNA.**

Table lists the name, strand, start, end, mean, and median enrichment factor of all common REZs in hmtDNA (see **Methods**), and all libraries containing these common REZs.

**Additional file 5 rNMP hotspots in all libraries and each subset in hmtDNA.**

Table represents rNMP hotspots in all libraries and each subset $CD4^{+}T$, DLTB, hESC-H9, HEK293T, and HEK293T RNH2A KO. The location, strand, and rNMP with dNMPs in +/- 3 nt locations of each hotspot are indicated in the table. Median EF represents the median value of the enrichment factor found in each library.

**Additional file 6 Statistical test for preferred rNMP composition and pattern.**

Mann-Whitney *U* test for rNMP composition and preferred dinucleotide and trinucleotide patterns in hmtDNA and *S. cerevisiae* mtDNA. Right-sided Mann-Whitney *U* tests are performed on the normalized frequencies versus the expectation value (0.25 for composition and dinucleotide pattern, 0.125 for trinucleotide pattern) for each cell type and all cell types combined. Cells with bold font show statistically significant results with a P-value < 0.05, which means the pattern is preferred.

**Additional file 7 Statistical test for rNMP-embedment composition and pattern bias between the two strands.**

Mann-Whitney *U* test for rNMP composition and preferred dinucleotide and trinucleotide patterns between light and heavy strands in hmtDNA and between forward and reverse strands in *S. cerevisiae.* Two-tailed paired Mann-Whitney *U* tests are performed on the normalized frequencies between both strands. Cells with bold font show statistically significant results with a P-value < 0.05, which means the pattern is biased between the two strands.

**Additional file 8 Maximum rNMP count in each library.**

The maximum rNMP counts at the single-nucleotide position on each strand are calculated. “Positive Maxcount” and “Negative Maxcount" in the table are the maximum rNMPs count in the light strand and heavy strand.

**Table S1. Oligonucleotides used in this study.**

The name, length, and sequence of oligonucleotides used in this study are presented. All bold letters in the PCR primers indicate the specific sequence of the index used in sequencing. P and Am indicate end modifications of phosphate and amino modifier, respectively. All oligonucleotides were desalted, except those marked with an asterisk (*), which were HPLC purified. All oligonucleotides were synthesized by Integrated DNA Technologies.

**Figure S1 Embedded rNMPs are prevalent on the reverse strand of *S. cerevisiae* mtDNA.**

(**A**) Bar graph showing the percentage of embedded rNMPs on the light (red bars) and heavy (blue bars) strands in yeast mtDNA. The green vertical line separates RNH201 WT and KO (*rnh201*) libraries. Two-tailed Mann-Whitney *U* test is performed to check the significance of the light/heavy strand bias. *: 0.01 < P < 0.05; **: 0.001 < P < 0.01; ***:

0.0001 < P < 0.001; ****: 0.00001 < P < 0.0001. (**B**) rNMP frequency at each nucleotide on the forward (top red spikes) and reverse (bottom blue spikes) strands of yeast mtDNA of ribose-seq (FS) and emRiboSeq (EM) libraries. Examples of consistent enrichment of rNMP on the light and heavy strands are indicated by red and blue arrows.

**Figure S2 rNMP distribution on both strands of hmtDNA.**
rNMP frequency at each nucleotide on the light (outer-circle red spikes) and heavy (inner-circle blue spikes) strands of hmtDNA in each library. The total numbers of rNMPs in each library are listed in **Additional file 1**.

**Figure S3 rNMP-enriched zones in the hmtDNA control region.**
rNMP embedment probability per base (PPB) in the hmtDNA control region for each cell category. The upper part of the figure is the rNMP PPB on the light strand and the lower one is on the heavy strand. The OriH region is shown in green color and the rest of the control region is shown in orange color. The standard deviation is indicated by the shadow.

**Figure S4 Embedded rNMPs inhibit the binding activity of Pol $\gamma$ and TWINKLE**
The rNMP PPB on the light and heavy strand of the control region in all RNH2A WT libraries with chromatin immunoprecipitation-sequencing (ChIP-seq) data of Pol $\gamma$ and TWINKLE (40). The blue and red bent arrows indicate the heavy-strand promoter (HSP) and light-strand promoter (LSP), respectively. Around the high PPB region, there is a lower ChIP-seq signal compared to neighborhood regions.

**Figure S5 The rNMP-embedment frequency does not correlate with CDS length on the template strand and non-template strand of non-coding genes.**
(**A**) rNMP-embedment probability per base (PPB) on the template strand of each coding sequence in each library. The PPB measures the normalized rNMP-embedment frequency at each base (see **Methods**). Each row represents a CDS, which is sorted by descending gene size. Each column represents an rNMP library. There are three control libraries with artificial rNMP embedment generated *in silico* on the right (see **Methods**). (**B**) Linear

regression plot showing no correlation between the rNMP enrichment factor and gene size on the template strand. Each data point represents the rNMP-enrichment factor on the template strand of a particular gene in an rNMP library of a given cell type. The rNMP-enrichment factors are calculated as described in Methods. Spearman's r and p-value are marked in the plot. Cell types with r > 0.5 have a strong positive correlation between the rNMP-enrichment factor and gene size. The 95% confidential interval is marked as the blue shadow region around the line. (**C**) Linear regression plot on both non-template and template strands for other cell categories. (**D**) rNTP-incorporation probability per base (PPB) on the non-template strand of each non-coding gene in each library.

**Figure S6 The rNMP-embedment frequency does not correlate with CDS length in *S. cerevisiae* mtDNA.**

rNMP-embedment probability per base (PPB) on the (**A**) non-template strand and (**B**) template strand of each coding sequence in each library of *S. cerevisiae* mtDNA. The PPB measures the normalized rNTP-incorporation frequency at each nucleotide (see **Methods**). Each row represents a gene, which is sorted by descending gene size. Each column represents an rNMP library.

**Figure S7 rNMP composition in mtDNA of each library.**

Heatmap analyses with the normalized frequency of incorporated rNMP (R: rA, rC, rG, or rU) on (**A**) both strands, (**B**) light strand, and (**C**) heavy strand of hmtDNA in all libraries.

**Figure S8. Consensus analysis for high-frequency rNMP location motifs in hmtDNA of the different cell types.**

Sequence-motif plots for high-frequency rNMP locations (Top 1% sites of most abundant embedded rNMPs, see **Methods**). Position 0 on the x-axis represents the rNMP site, - and + positions represent upstream and downstream dNMPs, respectively. The y-axis shows the level of sequence conservation, represented in bits. The cell type, genotype, library name, and the number of rNMP sites are included below each plot.

**Figure S9 rNMP composition in *S. cerevisiae*.**

(**A, B**) Bar plot showing the normalized frequency of the four rNMP types on the **(A)** forward strand and the **(B)** reverse strand of yeast mtDNA. The normalized frequency is calculated as described in the **Methods** section. Types of rNMPs and their corresponding colors are listed in the color legend on the right top. (**C, D**) Heatmap analyses with the normalized frequency of incorporated rNMP (R: rA, rC, rG, or rU) on the (**C**) forward strand and the (**D**) reverse strand of yeast mtDNA in all libraries. The vertical green line separates the cell types with RNH2A WT and with RNH2A KO genotype.

**Figure S10 Relative rNTP abundance in different human cell types.**

rNTP pool to dNTP pool ratios in total cells of $CD4^{+}T$, hESC-H9, HEK293T WT RNH2A, and RNH2A-KO. dNTP and rNTP concentration (pmol/million cells) for activated $CD4^{+}T$ cells were collected from a previous study having 4 replicates, and the precalculated average was used to represent dNTP/rNTP ratios in this figure (55). Data on the rest of the cell lines were collected for this study and nucleotide contents were quantified using Thermo TSQ Quantiva Lc-MS/MS with Kinetex EVO-C18 column. Pmol/million cells were calculated for rNTPs and dNTPs for all four replicates to calculate the ratios in this figure.

**Figure S11 Downstream dNMP is not associated with incorporated rNMP.**

**(A)** Heatmap analyses with the normalized frequency of dinucleotides composed of the incorporated rNMP (R: rA, rC, rG, or rU) and its downstream dNMP neighbor (N: dA, dC, dG, or dT) (RN) in the hmtDNA. The rNMP position in the dinucleotide is shown in red at the top left of the heatmap. Each column of the heatmap shows the results of an rNMP library. The vertical, green lines separate different cell types. The color scale is shown in the top right corner of the figure.

**Figure S12 Biased rNTP-incorporation dinucleotide patterns between the light and heavy strands of hmtDNA.**

Heatmap analyses with the normalized frequency of dinucleotides composed of the incorporated rNMP (R: rA, rC, rG, or rU) and its upstream dNMP neighbor (N: dA, dC, dG, or dT) (NR) in the (**A**) light and (**B**) heavy strand of hmtDNA. The rNMP position in the dinucleotide is shown in red at the top left of the heatmap. Each column of the heatmap shows the results of an rNMP library. The vertical, green lines separate different cell types. The color scale is shown in the top right corner of the figure.

**Figure S13 Similar rNTP-incorporation dinucleotide patterns between the forward and reverse strands in *S. cerevisiae* mtDNA.**

Heatmap analyses with the normalized frequency of dinucleotides composed of the incorporated rNMP (R: rA, rC, rG, or rU) and its upstream dNMP neighbor (N: dA, dC, dG, or dT) (NR) in the (**A**) forward and (**B**) reverse strand of yeast mtDNA. The rNMP position in the dinucleotide is shown in red at the top left of the heatmap. Each column of the heatmap shows the results of an rNMP library. The vertical, green lines separate different RNase H2 genotypes. The color scale is shown in the top right corner of the figure.

**Figure S14 Preferred trinucleotide of embedded rNMPs in hmtDNA.**

Heatmap analyses with the normalized frequency of trinucleotides composed of the incorporated rNMP (R: rA, rC, rG, or rU) and its two upstream dNMP neighbors (N: dA, dC, dG, or dT) (NNR) in the (**A**) both strands, (**B**) light strand, and (**C**) heavy strand of hmtDNA. The vertical, green lines separate different cell types. The color scale is shown in the bottom right corner of the figure.

**Figure S15 Preferred trinucleotide of embedded rNMPs in *S. cerevisiae* mtDNA.**

Heatmap analyses with the normalized frequency of trinucleotides composed of the incorporated rNMP (R: rA, rC, rG, or rU) and its two upstream dNMP neighbors (N: dA, dC, dG, or dT) (NNR) in the (**A**) both strands, (**B**) forward strand, and (**C**) reverse strand of yeast mtDNA. The vertical, green lines separate different RNase H2 genotypes. The color scale is shown in the bottom right corner of the figure.

**Table S1. Oligonucleotides used in this study.**

| Name | Size | Sequence |
|---|---|---|
| Adaptor.L1 * | 65 | 5’ P-NNC CGN NNN NNA GAT CGG AAG AGC GTC GTG TAG GGA AAG AGT GTT GAT AGA TCC GTG TCG CAA C*T |
| Adaptor.L2 * | 65 | 5’ P-NNT GAN NNN NNA GAT CGG AAG AGC GTC GTG TAG GGA AAG AGT GTT GAT AGA TCC GTG TCG CAA C*T |
| Adaptor.L3 * | 65 | 5’ P-NNG ACN NNN NNA GAT CGG AAG AGC GTC GTG TAG GGA AAG AGT GTT GAT AGA TCC GTG TCG CAA C*T |
| Adaptor.L4 * | 65 | 5’ P-NNG AAN NNN NNA GAT CGG AAG AGC GTC GTG TAG GGA AAG AGT GTT GAT AGA TCC GTG TCG CAA C*T |
| Adaptor.L5 * | 65 | 5’ P-NNG CTN NNN NNA GAT CGG AAG AGC GTC GTG TAG GGA AAG AGT GTT GAT AGA TCC GTG TCG CAA C*T |
| Adaptor.L6 * | 65 | 5’ P-NNA GCN NNN NNA GAT CGG AAG AGC GTC GTG TAG GGA AAG AGT GTT GAT AGA TCC GTG TCG CAA C*T |
| Adaptor.L7 * | 65 | 5’ P-NNC TGN NNN NNA GAT CGG AAG AGC GTC GTG TAG GGA AAG AGT GTT GAT AGA TCC GTG TCG CAA C*T |
| Adaptor.L8 * | 65 | 5’ P-NNT GTN NNN NNA GAT CGG AAG AGC GTC GTG TAG GGA AAG AGT GTT GAT AGA TCC GTG TCG CAA C*T |
| Adaptor.S * | 25 | 5’ P-GTT GCG ACA CGG ATC TAT CAA CAC T -Am 3’ |
| PCR.1 | 54 | 5’ GTG ACT GGA GTT CAG ACG TGT GCT CTT CCG ATC TTG ATA GAT CCG TGT CGC AAC |
| PCR.2 | 20 | 5’ ACA CTC TTT CCC TAC ACG AC |
| PCR.701 | 53 | 5’ CAA GCA GAA GAC GGC ATA CGA GAT **CGA GTA AT**G TGA CTG GAG TTC AGA CGT GT |
| PCR.702 | 53 | 5’ CAA GCA GAA GAC GGC ATA CGA GAT **TCT CCG GA**G TGA CTG GAG TTC AGA CGT GT |
| PCR.705 | 53 | 5' CAA GCA GAA GAC GGC ATA CGA GAT **TTC TGA AT**G TGA CTG GAG TTC AGA CGT GT |
| PCR.707 | 53 | 5' CAA GCA GAA GAC GGC ATA CGA GAT **AGC TTC AG**G TGA CTG GAG TTC AGA CGT GT |

**Table S1 continued**

| | | |
|---|---|---|
| PCR.712 | 53 | 5' CAA GCA GAA GAC GGC ATA CGA GAT **CTA TCG CT**G TGA CTG GAG TTC AGA CGT GT |
| PCR.501 | 57 | 5’ AAT GAT ACG GCG ACC GAG ATC TAC AC**T ATA GCC T**AC ACT CTT TCC CTA CAC GAC |
| PCR.502 | 57 | 5’ AAT GAT ACG GCG ACC GAG ATC TAC AC**A TAG AGG C**AC ACT CTT TCC CTA CAC GAC |
| PCR.503 | 57 | 5’ AAT GAT ACG GCG ACC GAG ATC TAC AC**C CTA TCC T**AC ACT CTT TCC CTA CAC GAC |
| PCR.504 | 57 | 5’ AAT GAT ACG GCG ACC GAG ATC TAC AC**G GCT CTG A**AC ACT CTT TCC CTA CAC GAC |
| PCR.505 | 57 | 5’ AAT GAT ACG GCG ACC GAG ATC TAC AC**A GGC GAA G**AC ACT CTT TCC CTA CAC GAC |
| PCR.506 | 57 | 5’ AAT GAT ACG GCG ACC GAG ATC TAC AC**T AAT CTT A**AC ACT CTT TCC CTA CAC GAC |
| PCR.507 | 57 | 5’ AAT GAT ACG GCG ACC GAG ATC TAC AC**C AGG ACG T**AC ACT CTT TCC CTA CAC GAC |
| PCR.508 | 57 | 5’ AAT GAT ACG GCG ACC GAG ATC TAC AC**G TAC TGA C**AC ACT CTT TCC CTA CAC GAC |

Figure S1

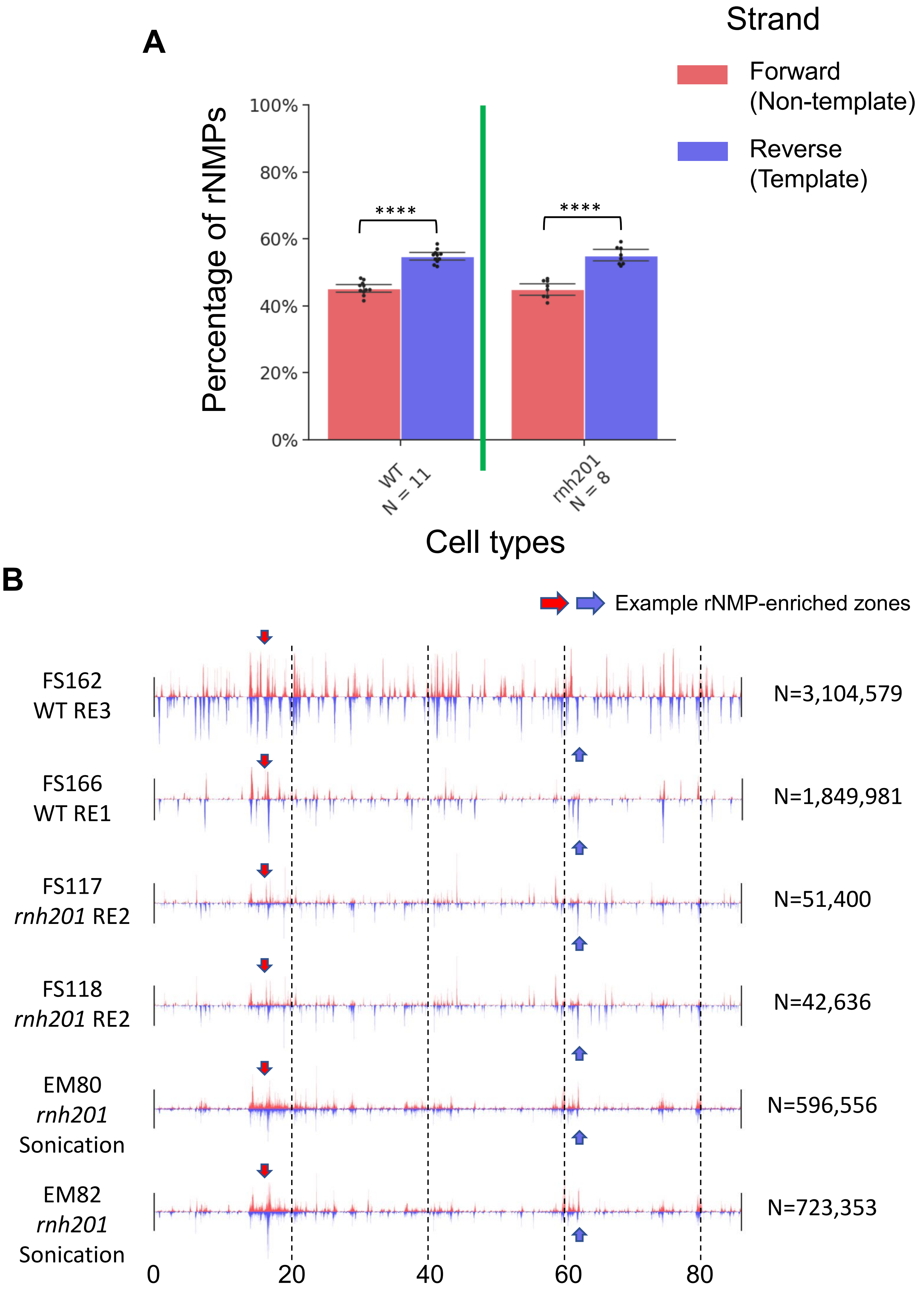
A
Strand
Forward
(Non-template)
Reverse
(Template)
Percentage of rNMPs
100%
80%
60%
40%
20%
0%
****
****
WT
N = 11
rnh201
N = 8
Cell types
B
Example rNMP-enriched zones
FS162
WT RE3
N=3,104,579
FS166
WT RE1
N=1,849,981
FS117
rnh201 RE2
N=51,400
FS118
rnh201 RE2
N=42,636
EM80
rnh201
Sonication
N=596,556
EM82
rnh201
Sonication
N=723,353
0
20
40
60
80

### Figure S2

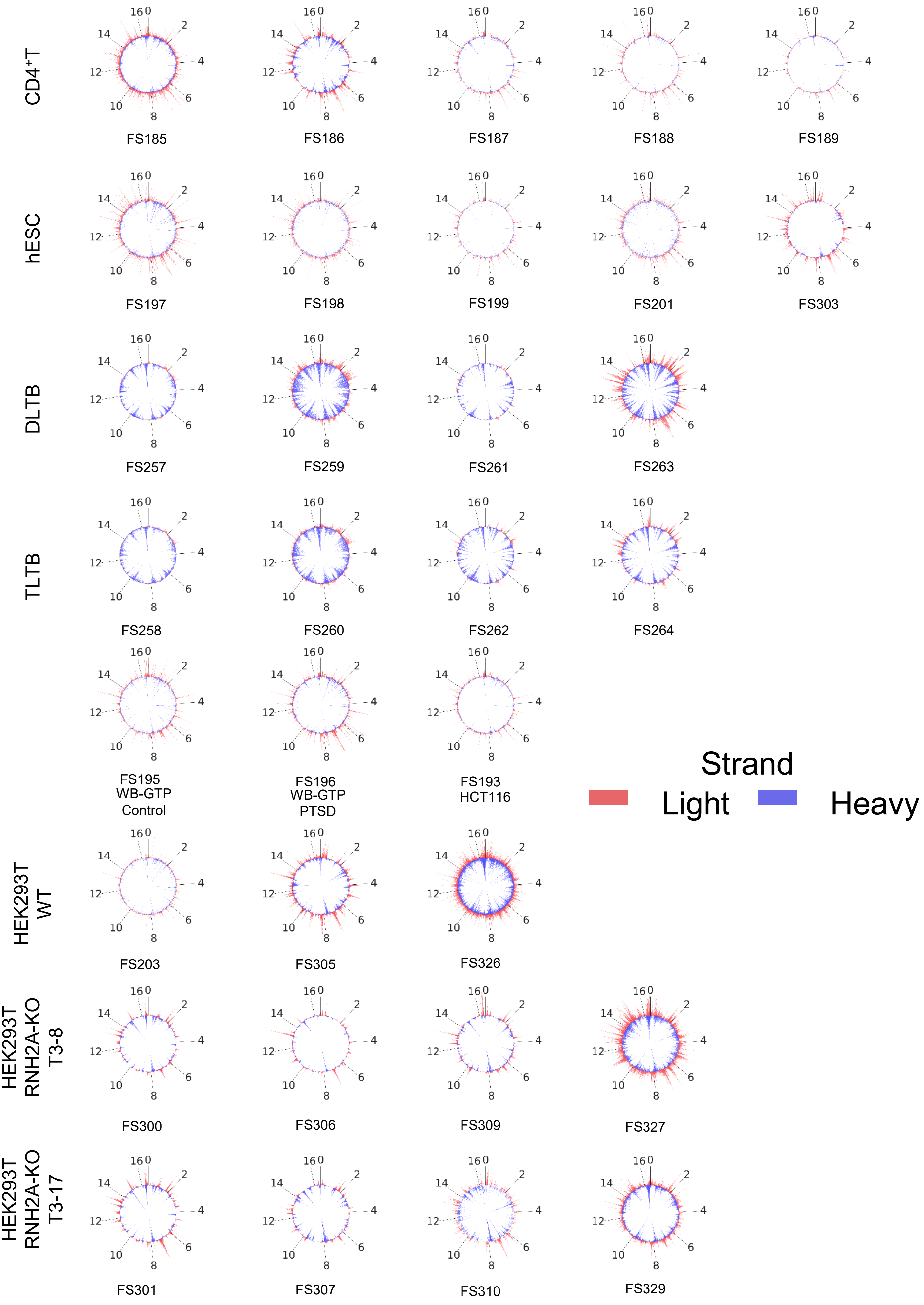
CD4+T
FS185
FS186
FS187
FS188
FS189
hESC
FS197
FS198
FS199
FS201
FS303
DLTB
FS257
FS259
FS261
FS263
TLTB
FS258
FS260
FS262
FS264
FS195
WB-GTP
Control
FS196
WB-GTP
PTSD
FS193
HCT116
Strand
Light
Heavy
HEK293T
WT
FS203
FS305
FS326
HEK293T
RNH2A-KO
T3-8
FS300
FS306
FS309
FS327
HEK293T
RNH2A-KO
T3-17
FS301
FS307
FS310
FS329

Figure S3

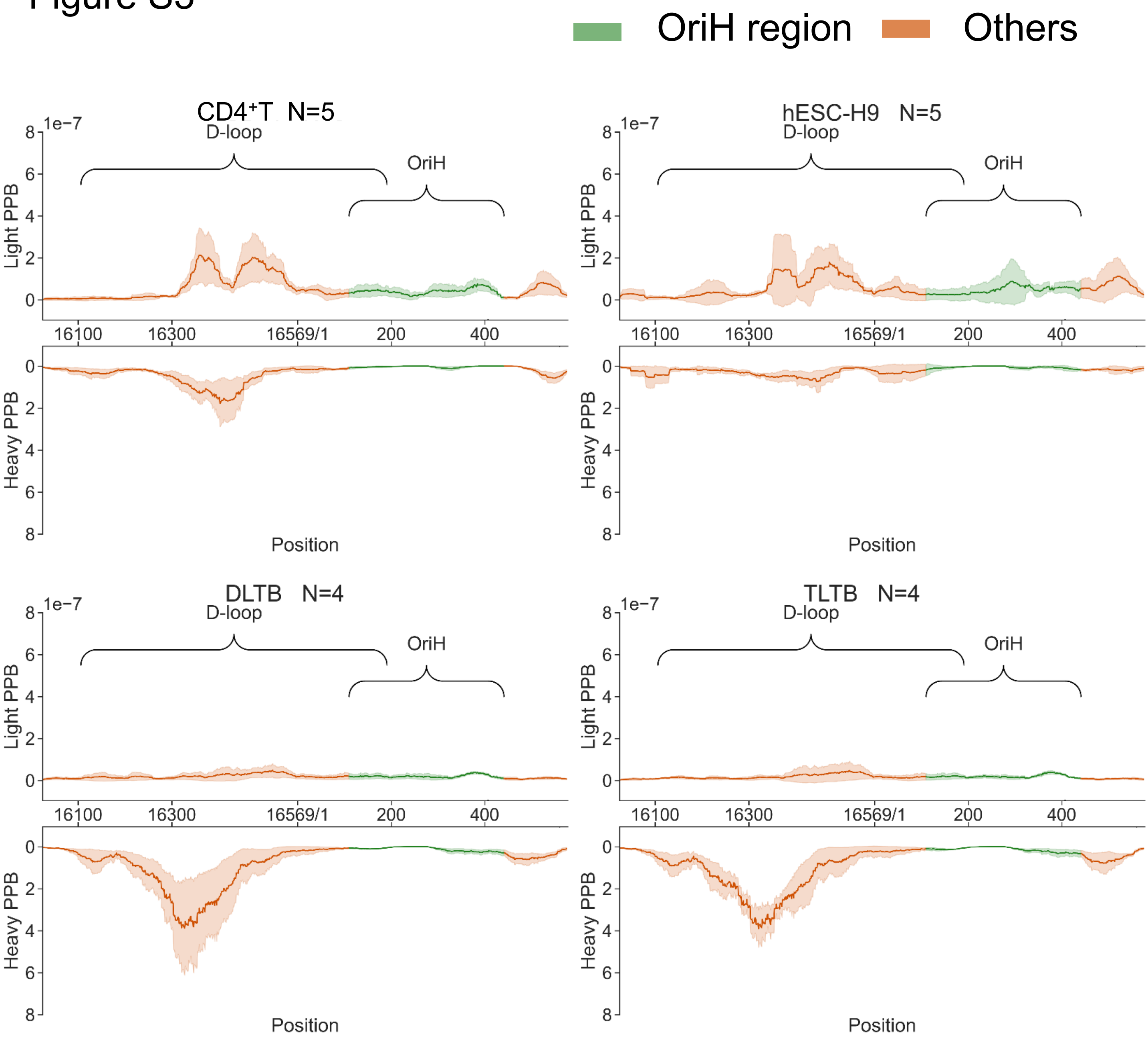
OriH region
Others
CD4⁺T N=5
hESC-H9 N=5
DLTB N=4
TLTB N=4
1e−7
D-loop
OriH
Light PPB
Heavy PPB
0
2
4
6
8
16100
16300
16569/1
200
400
Position

### Figure S3 continued

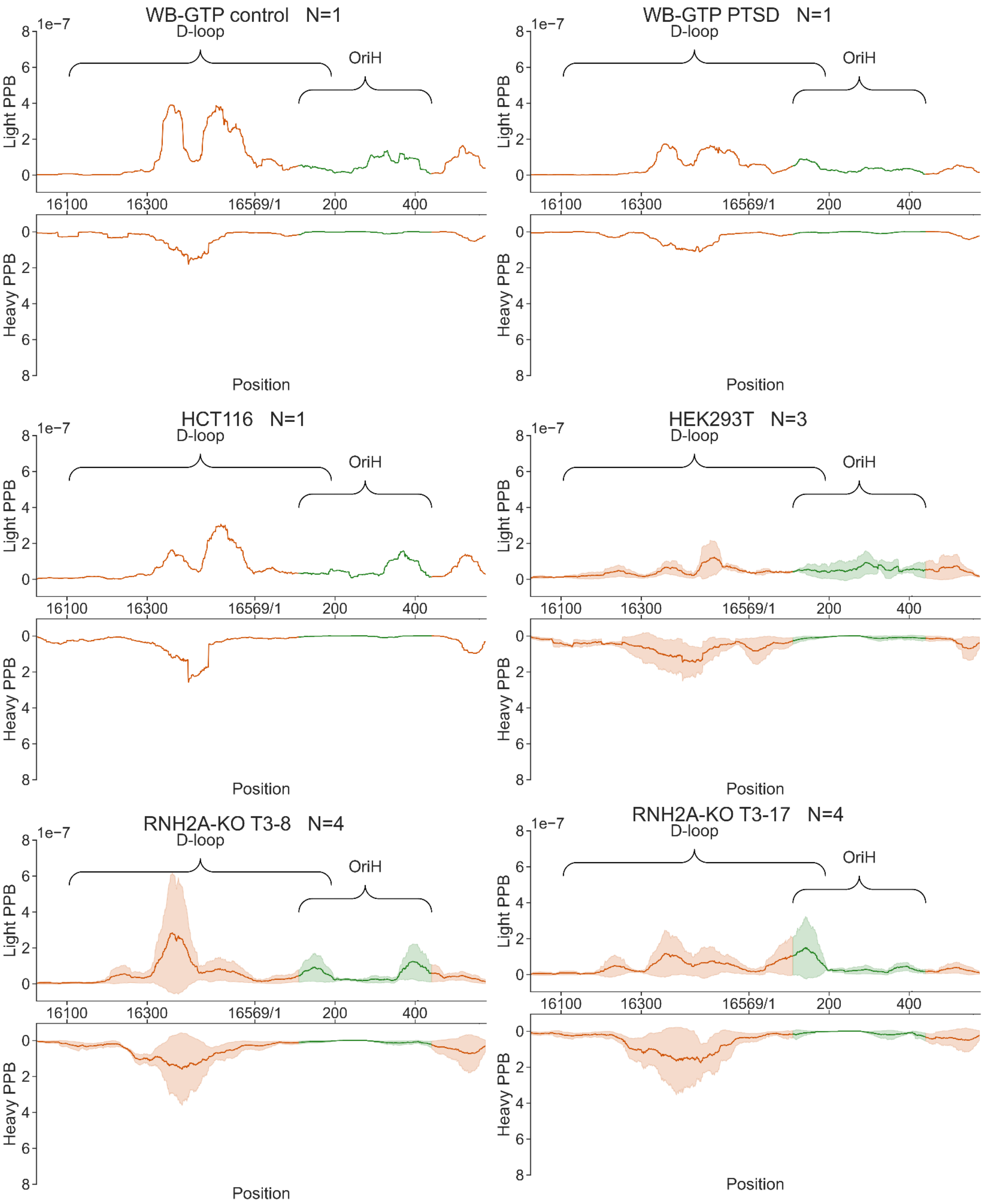

### Figure S4

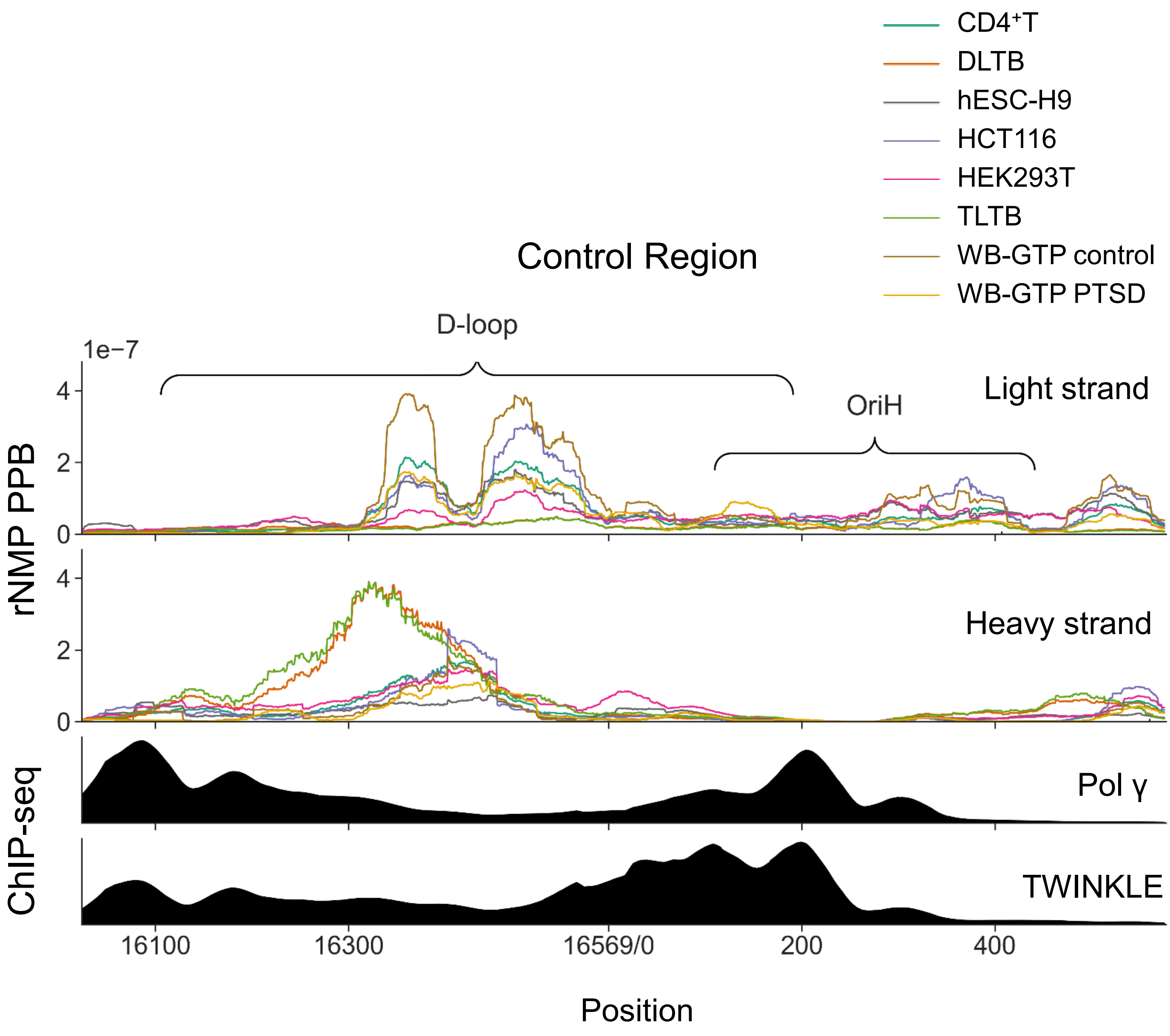

CD4⁺T
DLTB
hESC-H9
HCT116
HEK293T
TLTB
WB-GTP control
WB-GTP PTSD
Control Region
D-loop
1e−7
OriH
Light strand
Heavy strand
rNMP PPB
4
2
0
4
2
0
Pol γ
TWINKLE
ChIP-seq
16100
16300
16569/0
200
400
Position

Figure S5

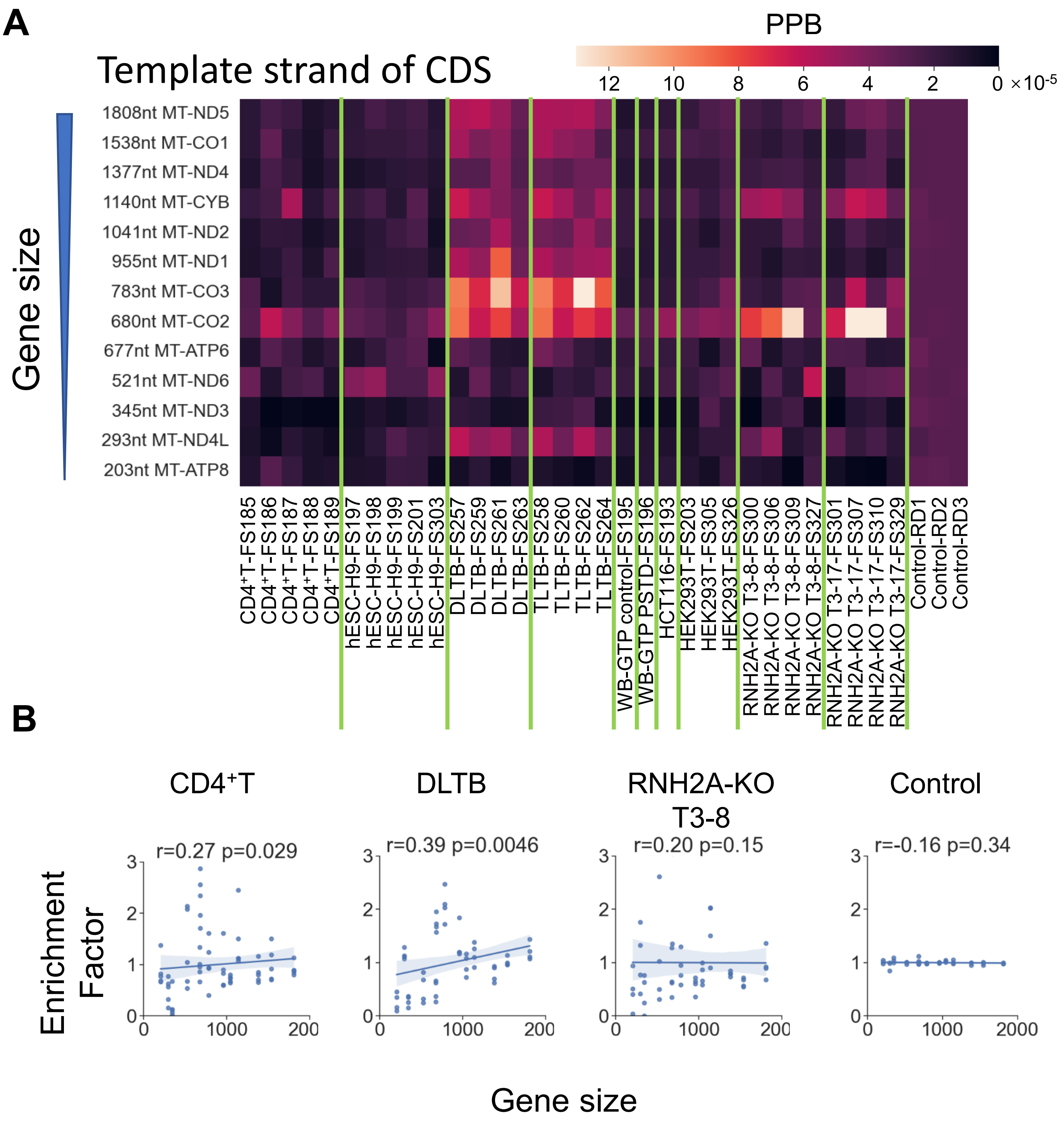
A
Template strand of CDS
PPB
12 10 8 6 4 2 0 ×10-5
Gene size
1808nt MT-ND5
1538nt MT-CO1
1377nt MT-ND4
1140nt MT-CYB
1041nt MT-ND2
955nt MT-ND1
783nt MT-CO3
680nt MT-CO2
677nt MT-ATP6
521nt MT-ND6
345nt MT-ND3
293nt MT-ND4L
203nt MT-ATP8
CD4+T-FS185
CD4+T-FS186
CD4+T-FS187
CD4+T-FS188
CD4+T-FS189
hESC-H9-FS197
hESC-H9-FS198
hESC-H9-FS199
hESC-H9-FS201
hESC-H9-FS303
DLTB-FS257
DLTB-FS259
DLTB-FS261
DLTB-FS263
TLTB-FS258
TLTB-FS260
TLTB-FS262
TLTB-FS264
WB-GTP control-FS195
WB-GTP PSTD-FS196
HCT116-FS193
HEK293T-FS203
HEK293T-FS305
HEK293T-FS326
RNH2A-KO T3-8-FS300
RNH2A-KO T3-8-FS306
RNH2A-KO T3-8-FS309
RNH2A-KO T3-8-FS327
RNH2A-KO T3-17-FS301
RNH2A-KO T3-17-FS307
RNH2A-KO T3-17-FS310
RNH2A-KO T3-17-FS329
Control-RD1
Control-RD2
Control-RD3
B
CD4+T
r=0.27 p=0.029
DLTB
r=0.39 p=0.0046
RNH2A-KO T3-8
r=0.20 p=0.15
Control
r=-0.16 p=0.34
Enrichment Factor
Gene size
0 1 2 3
0 1000 2000

Figure S5 continued

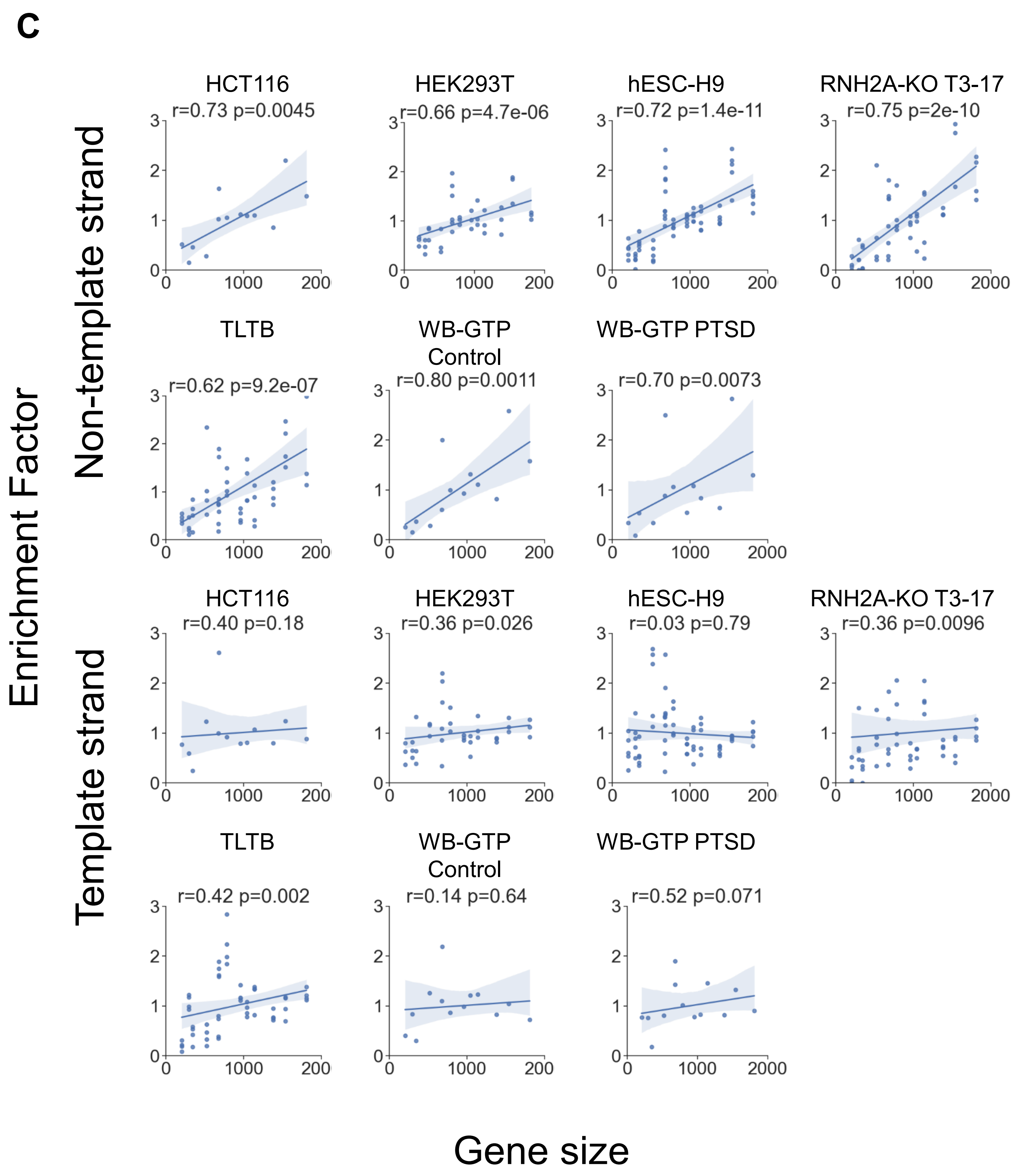

Figure S5 continued

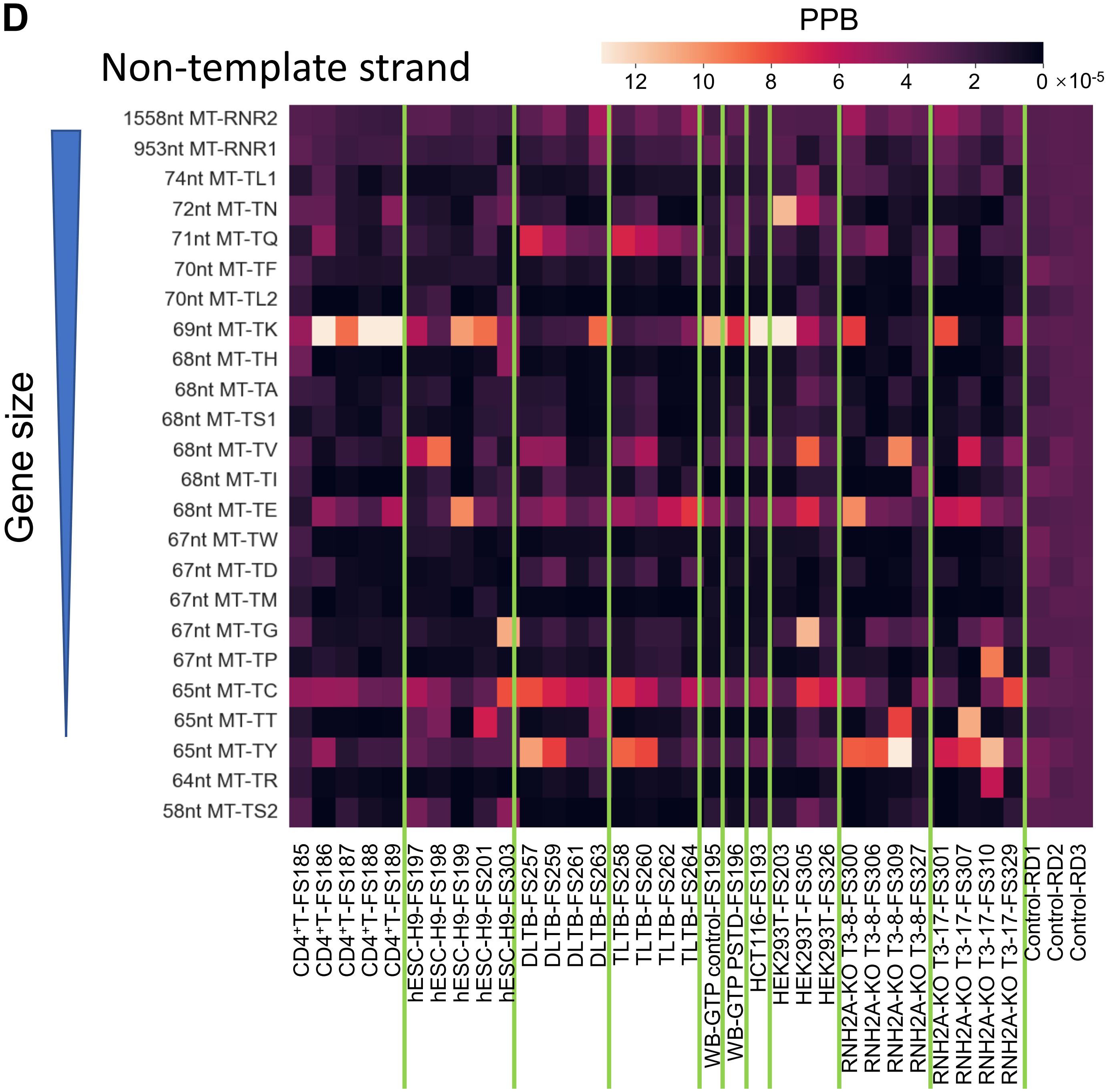

Figure S6

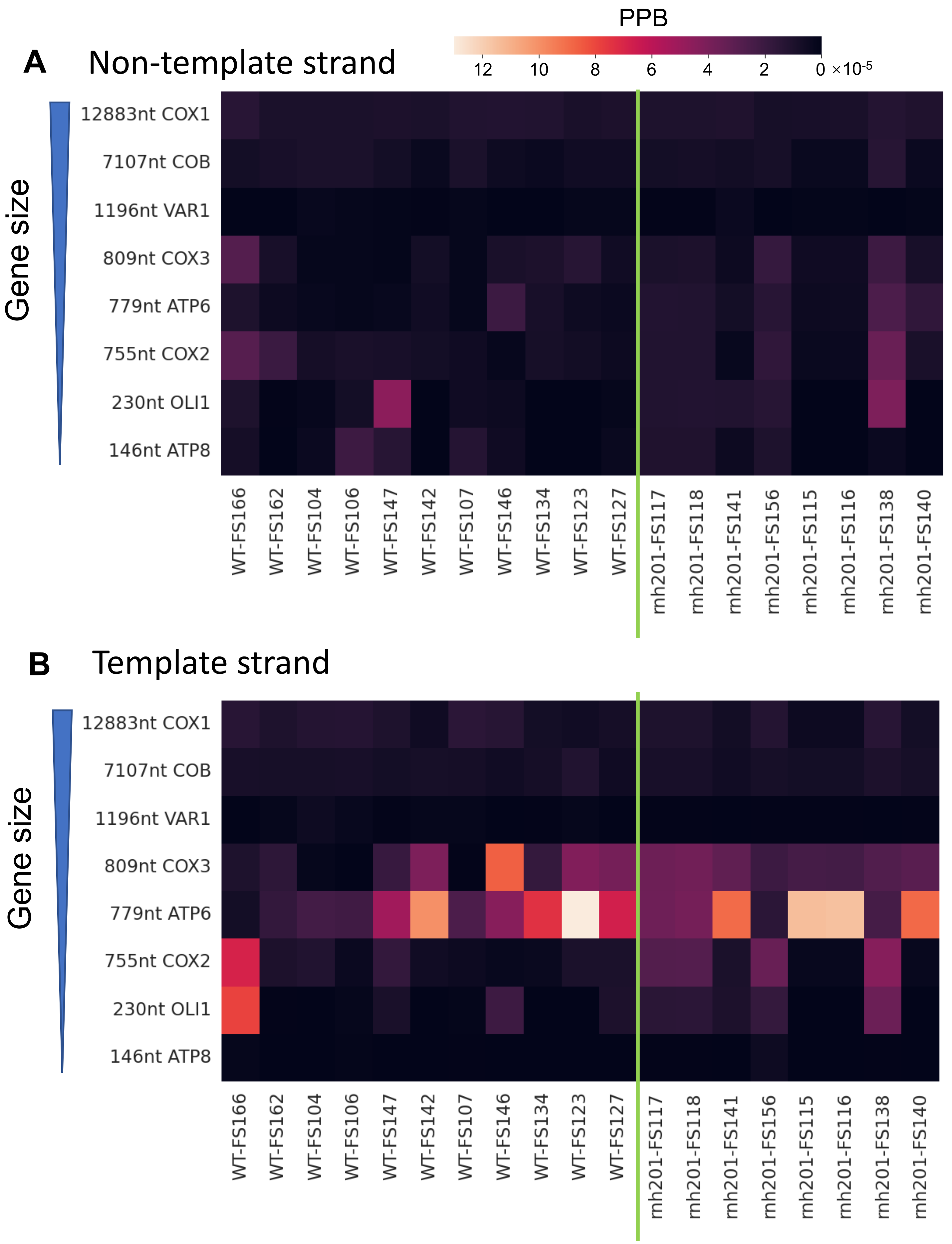

PPB
12
10
8
6
4
2
0 ×10-5
A
Non-template strand
Gene size
12883nt COX1
7107nt COB
1196nt VAR1
809nt COX3
779nt ATP6
755nt COX2
230nt OLI1
146nt ATP8
WT-FS166
WT-FS162
WT-FS104
WT-FS106
WT-FS147
WT-FS142
WT-FS107
WT-FS146
WT-FS134
WT-FS123
WT-FS127
mh201-FS117
mh201-FS118
mh201-FS141
mh201-FS156
mh201-FS115
mh201-FS116
mh201-FS138
mh201-FS140
B
Template strand
Gene size
12883nt COX1
7107nt COB
1196nt VAR1
809nt COX3
779nt ATP6
755nt COX2
230nt OLI1
146nt ATP8
WT-FS166
WT-FS162
WT-FS104
WT-FS106
WT-FS147
WT-FS142
WT-FS107
WT-FS146
WT-FS134
WT-FS123
WT-FS127
mh201-FS117
mh201-FS118
mh201-FS141
mh201-FS156
mh201-FS115
mh201-FS116
mh201-FS138
mh201-FS140

Figure S7

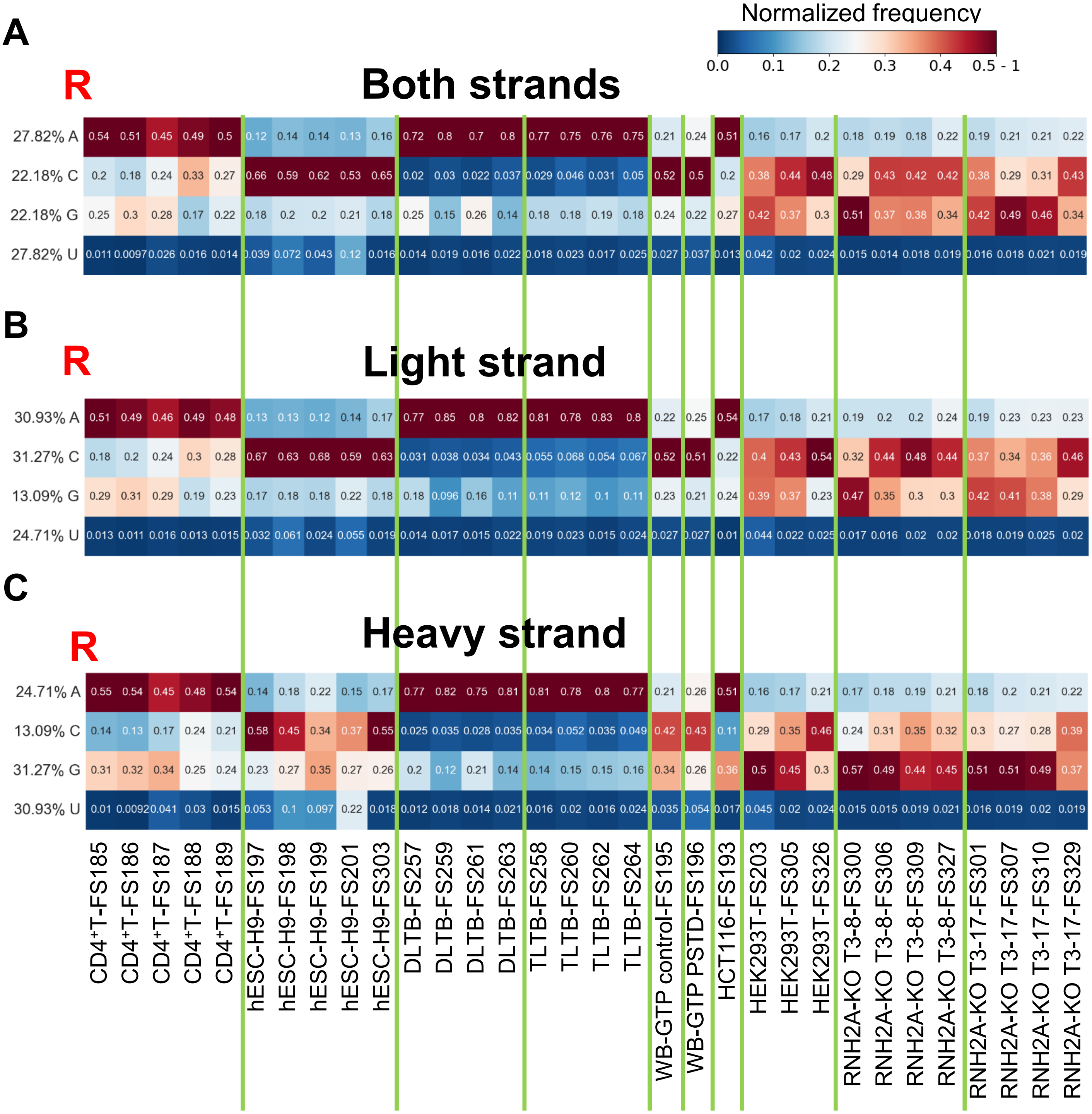
Normalized frequency
0.0 0.1 0.2 0.3 0.4 0.5 - 1
A
R
Both strands
27.82% A 0.54 0.51 0.45 0.49 0.5 0.12 0.14 0.14 0.13 0.16 0.72 0.8 0.7 0.8 0.77 0.75 0.76 0.75 0.21 0.24 0.51 0.16 0.17 0.2 0.18 0.19 0.18 0.22 0.19 0.21 0.21 0.22
22.18% C 0.2 0.18 0.24 0.33 0.27 0.66 0.59 0.62 0.53 0.65 0.02 0.03 0.022 0.037 0.029 0.046 0.031 0.05 0.52 0.5 0.2 0.38 0.44 0.48 0.29 0.43 0.42 0.42 0.38 0.29 0.31 0.43
22.18% G 0.25 0.3 0.28 0.17 0.22 0.18 0.2 0.2 0.21 0.18 0.25 0.15 0.26 0.14 0.18 0.18 0.19 0.18 0.24 0.22 0.27 0.42 0.37 0.3 0.51 0.37 0.38 0.34 0.42 0.49 0.46 0.34
27.82% U 0.011 0.0097 0.026 0.016 0.014 0.039 0.072 0.043 0.12 0.016 0.014 0.019 0.016 0.022 0.018 0.023 0.017 0.025 0.027 0.037 0.013 0.042 0.02 0.024 0.015 0.014 0.018 0.019 0.016 0.018 0.021 0.019
B
R
Light strand
30.93% A 0.51 0.49 0.46 0.49 0.48 0.13 0.13 0.12 0.14 0.17 0.77 0.85 0.8 0.82 0.81 0.78 0.83 0.8 0.22 0.25 0.54 0.17 0.18 0.21 0.19 0.2 0.2 0.24 0.19 0.23 0.23 0.23
31.27% C 0.18 0.2 0.24 0.3 0.28 0.67 0.63 0.68 0.59 0.63 0.031 0.038 0.034 0.043 0.055 0.068 0.054 0.067 0.52 0.51 0.22 0.4 0.43 0.54 0.32 0.44 0.48 0.44 0.37 0.34 0.36 0.46
13.09% G 0.29 0.31 0.29 0.19 0.23 0.17 0.18 0.18 0.22 0.18 0.18 0.096 0.16 0.11 0.11 0.12 0.1 0.11 0.23 0.21 0.24 0.39 0.37 0.23 0.47 0.35 0.3 0.3 0.42 0.41 0.38 0.29
24.71% U 0.013 0.011 0.016 0.013 0.015 0.032 0.061 0.024 0.055 0.019 0.014 0.017 0.015 0.022 0.019 0.023 0.015 0.024 0.027 0.027 0.01 0.044 0.022 0.025 0.017 0.016 0.02 0.02 0.018 0.019 0.025 0.02
C
R
Heavy strand
24.71% A 0.55 0.54 0.45 0.48 0.54 0.14 0.18 0.22 0.15 0.17 0.77 0.82 0.75 0.81 0.81 0.78 0.8 0.77 0.21 0.26 0.51 0.16 0.17 0.21 0.17 0.18 0.19 0.21 0.18 0.2 0.21 0.22
13.09% C 0.14 0.13 0.17 0.24 0.21 0.58 0.45 0.34 0.37 0.55 0.025 0.035 0.028 0.035 0.034 0.052 0.035 0.049 0.42 0.43 0.11 0.29 0.35 0.46 0.24 0.31 0.35 0.32 0.3 0.27 0.28 0.39
31.27% G 0.31 0.32 0.34 0.25 0.24 0.23 0.27 0.35 0.27 0.26 0.2 0.12 0.21 0.14 0.14 0.15 0.15 0.16 0.34 0.26 0.36 0.5 0.45 0.3 0.57 0.49 0.44 0.45 0.51 0.51 0.49 0.37
30.93% U 0.01 0.0092 0.041 0.03 0.015 0.053 0.1 0.097 0.22 0.018 0.012 0.018 0.014 0.021 0.016 0.02 0.016 0.024 0.035 0.054 0.017 0.045 0.02 0.024 0.015 0.015 0.019 0.021 0.016 0.019 0.02 0.019
CD4+T-FS185
CD4+T-FS186
CD4+T-FS187
CD4+T-FS188
CD4+T-FS189
hESC-H9-FS197
hESC-H9-FS198
hESC-H9-FS199
hESC-H9-FS201
hESC-H9-FS303
DLTB-FS257
DLTB-FS259
DLTB-FS261
DLTB-FS263
TLTB-FS258
TLTB-FS260
TLTB-FS262
TLTB-FS264
WB-GTP control-FS195
WB-GTP PSTD-FS196
HCT116-FS193
HEK293T-FS203
HEK293T-FS305
HEK293T-FS326
RNH2A-KO T3-8-FS300
RNH2A-KO T3-8-FS306
RNH2A-KO T3-8-FS309
RNH2A-KO T3-8-FS327
RNH2A-KO T3-17-FS301
RNH2A-KO T3-17-FS307
RNH2A-KO T3-17-FS310
RNH2A-KO T3-17-FS329

Figure S8

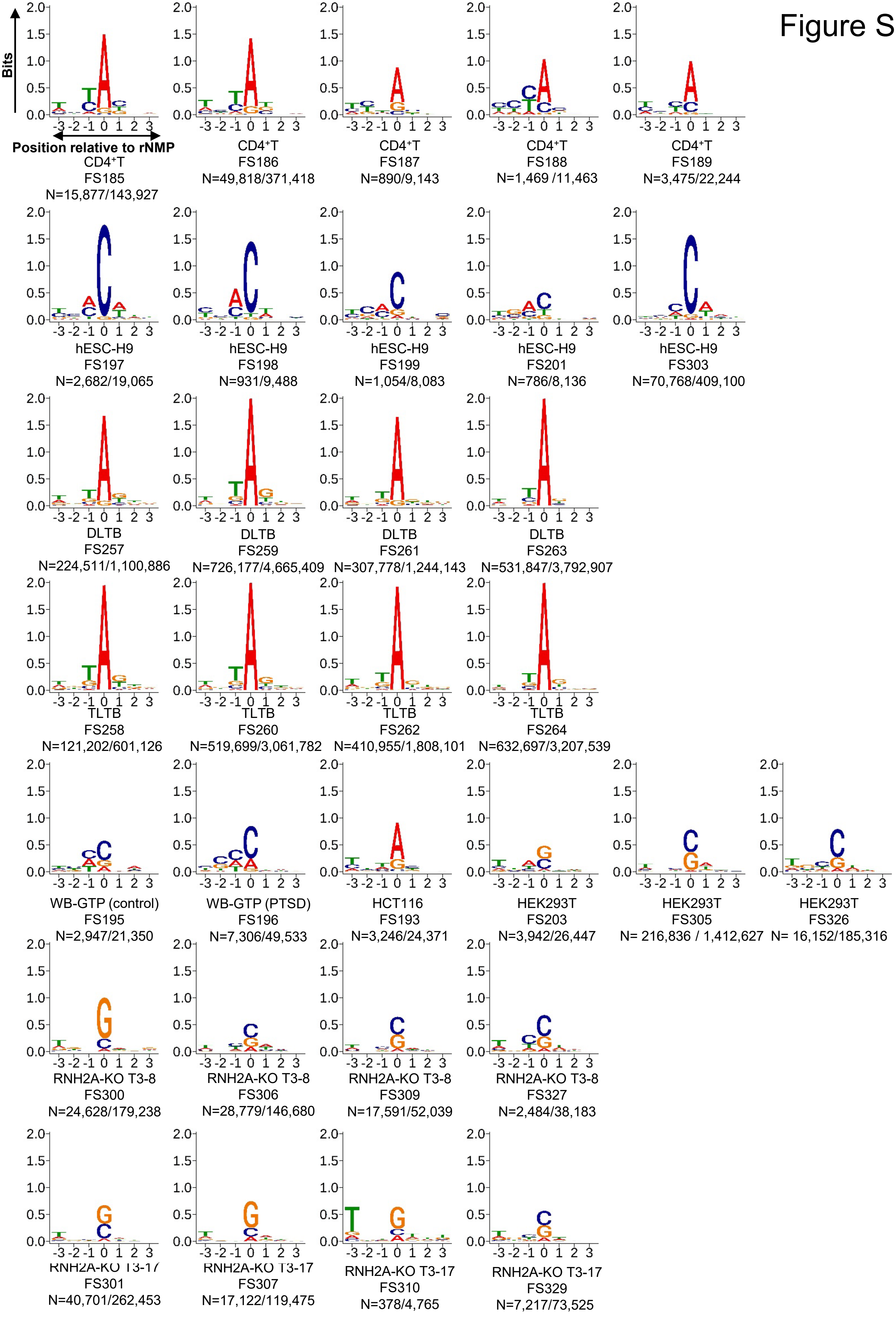
Bits
Position relative to rNMP
CD4⁺T FS185 N=15,877/143,927
CD4⁺T FS186 N=49,818/371,418
CD4⁺T FS187 N=890/9,143
CD4⁺T FS188 N=1,469 /11,463
CD4⁺T FS189 N=3,475/22,244
hESC-H9 FS197 N=2,682/19,065
hESC-H9 FS198 N=931/9,488
hESC-H9 FS199 N=1,054/8,083
hESC-H9 FS201 N=786/8,136
hESC-H9 FS303 N=70,768/409,100
DLTB FS257 N=224,511/1,100,886
DLTB FS259 N=726,177/4,665,409
DLTB FS261 N=307,778/1,244,143
DLTB FS263 N=531,847/3,792,907
TLTB FS258 N=121,202/601,126
TLTB FS260 N=519,699/3,061,782
TLTB FS262 N=410,955/1,808,101
TLTB FS264 N=632,697/3,207,539
WB-GTP (control) FS195 N=2,947/21,350
WB-GTP (PTSD) FS196 N=7,306/49,533
HCT116 FS193 N=3,246/24,371
HEK293T FS203 N=3,942/26,447
HEK293T FS305 N= 216,836 / 1,412,627
HEK293T FS326 N= 16,152/185,316
RNH2A-KO T3-8 FS300 N=24,628/179,238
RNH2A-KO T3-8 FS306 N=28,779/146,680
RNH2A-KO T3-8 FS309 N=17,591/52,039
RNH2A-KO T3-8 FS327 N=2,484/38,183
RNH2A-KO T3-17 FS301 N=40,701/262,453
RNH2A-KO T3-17 FS307 N=17,122/119,475
RNH2A-KO T3-17 FS310 N=378/4,765
RNH2A-KO T3-17 FS329 N=7,217/73,525

Figure S9

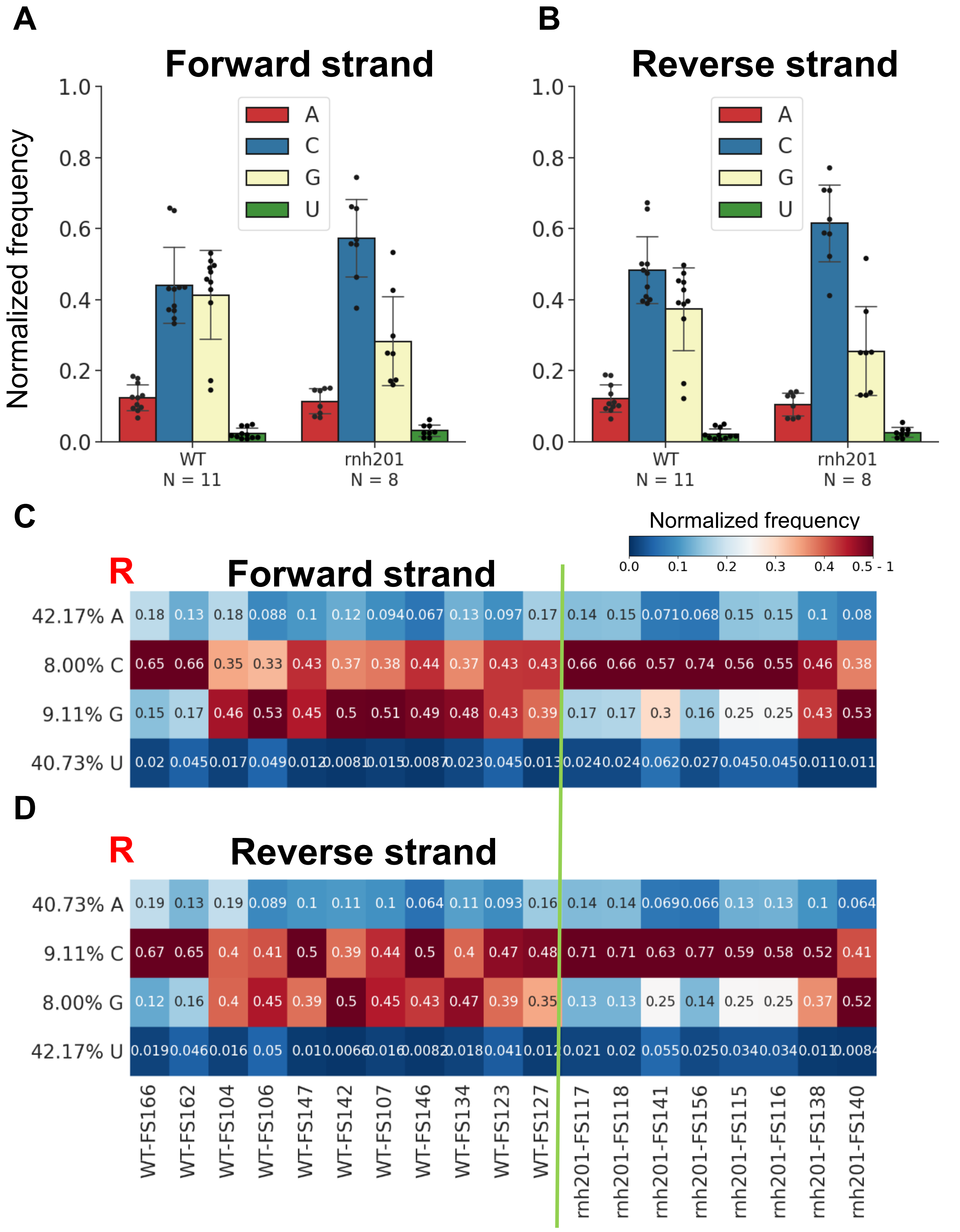

A
Forward strand
Normalized frequency
1.0
0.8
0.6
0.4
0.2
0.0
A
C
G
U
WT
N = 11
rnh201
N = 8
B
Reverse strand
1.0
0.8
0.6
0.4
0.2
0.0
A
C
G
U
WT
N = 11
rnh201
N = 8
C
R
Forward strand
Normalized frequency
0.0 0.1 0.2 0.3 0.4 0.5 - 1
42.17% A
8.00% C
9.11% G
40.73% U
0.18 0.13 0.18 0.088 0.1 0.12 0.094 0.067 0.13 0.097 0.17 0.14 0.15 0.071 0.068 0.15 0.15 0.1 0.08
0.65 0.66 0.35 0.33 0.43 0.37 0.38 0.44 0.37 0.43 0.43 0.66 0.66 0.57 0.74 0.56 0.55 0.46 0.38
0.15 0.17 0.46 0.53 0.45 0.5 0.51 0.49 0.48 0.43 0.39 0.17 0.17 0.3 0.16 0.25 0.25 0.43 0.53
0.02 0.045 0.017 0.049 0.012 0.0081 0.015 0.0087 0.023 0.045 0.013 0.024 0.024 0.062 0.027 0.045 0.045 0.011 0.011
D
R
Reverse strand
40.73% A
9.11% C
8.00% G
42.17% U
0.19 0.13 0.19 0.089 0.1 0.11 0.1 0.064 0.11 0.093 0.16 0.14 0.14 0.069 0.066 0.13 0.13 0.1 0.064
0.67 0.65 0.4 0.41 0.5 0.39 0.44 0.5 0.4 0.47 0.48 0.71 0.71 0.63 0.77 0.59 0.58 0.52 0.41
0.12 0.16 0.4 0.45 0.39 0.5 0.45 0.43 0.47 0.39 0.35 0.13 0.13 0.25 0.14 0.25 0.25 0.37 0.52
0.019 0.046 0.016 0.05 0.01 0.0066 0.016 0.0082 0.018 0.041 0.012 0.021 0.02 0.055 0.025 0.034 0.034 0.011 0.0084
WT-FS166
WT-FS162
WT-FS104
WT-FS106
WT-FS147
WT-FS142
WT-FS107
WT-FS146
WT-FS134
WT-FS123
WT-FS127
rnh201-FS117
rnh201-FS118
rnh201-FS141
rnh201-FS156
rnh201-FS115
rnh201-FS116
rnh201-FS138
rnh201-FS140

Figure S10

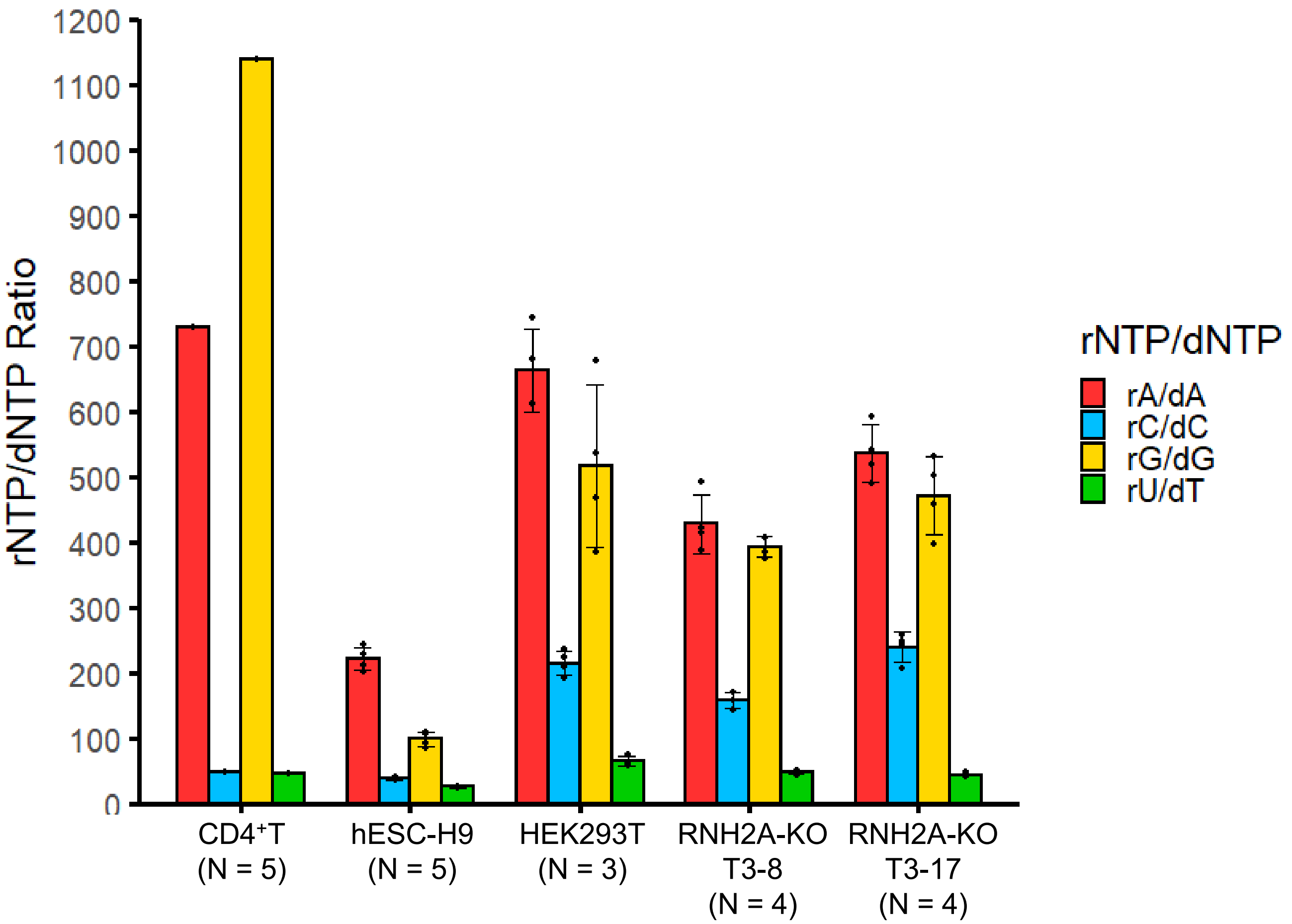

rNTP/dNTP Ratio
1200
1100
1000
900
800
700
600
500
400
300
200
100
0
CD4⁺T (N = 5)
hESC-H9 (N = 5)
HEK293T (N = 3)
RNH2A-KO T3-8 (N = 4)
RNH2A-KO T3-17 (N = 4)
rNTP/dNTP
rA/dA
rC/dC
rG/dG
rU/dT

Figure S11

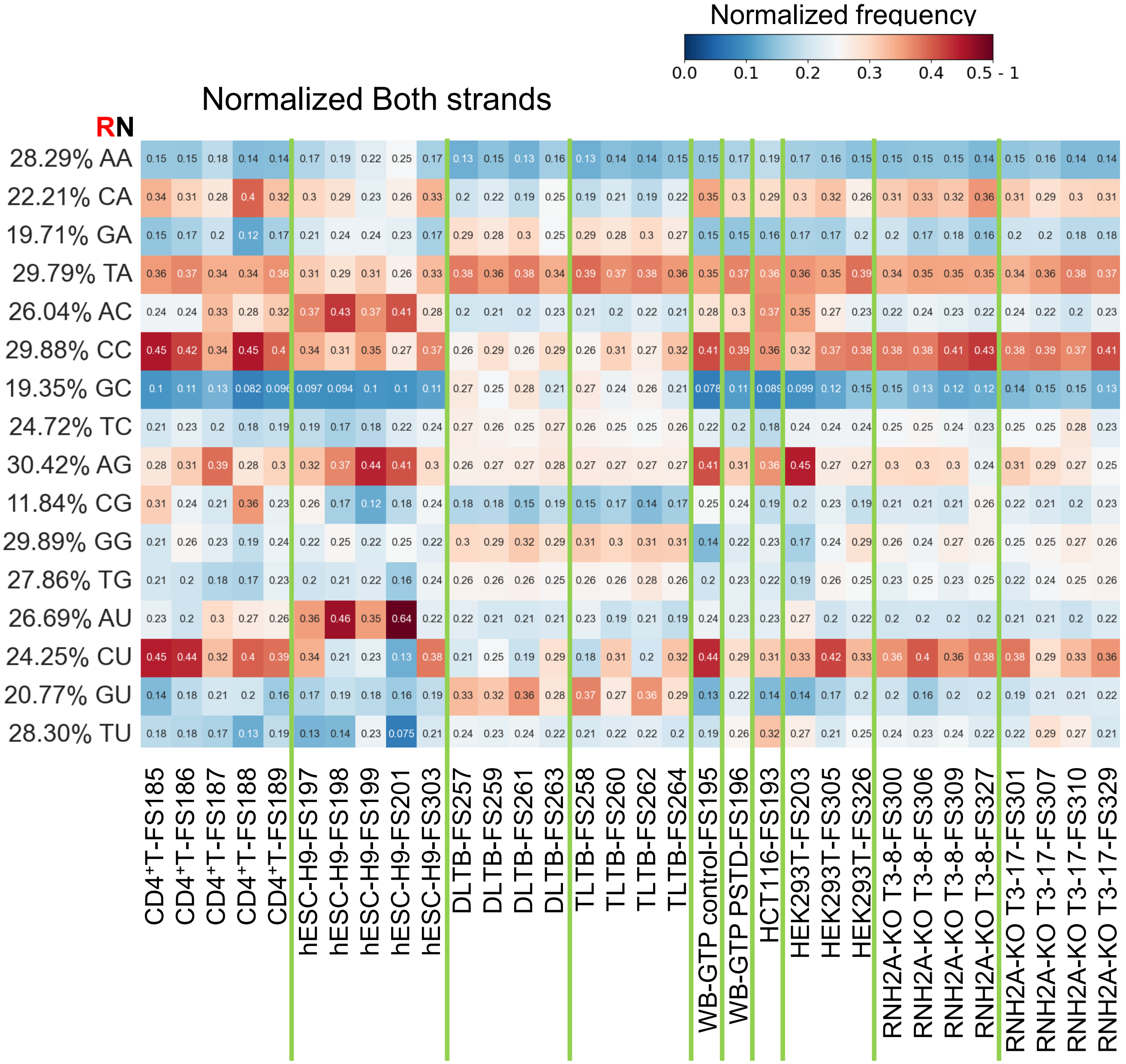
Normalized frequency
0.0 0.1 0.2 0.3 0.4 0.5 - 1
Normalized Both strands
RN
28.29% AA 0.15 0.15 0.18 0.14 0.14 0.17 0.19 0.22 0.25 0.17 0.13 0.15 0.13 0.16 0.13 0.14 0.14 0.15 0.15 0.17 0.19 0.17 0.16 0.15 0.15 0.15 0.15 0.14 0.15 0.16 0.14 0.14
22.21% CA 0.34 0.31 0.28 0.4 0.32 0.3 0.29 0.23 0.26 0.33 0.2 0.22 0.19 0.25 0.19 0.21 0.19 0.22 0.35 0.3 0.29 0.3 0.32 0.26 0.31 0.33 0.32 0.36 0.31 0.29 0.3 0.31
19.71% GA 0.15 0.17 0.2 0.12 0.17 0.21 0.24 0.24 0.23 0.17 0.29 0.28 0.3 0.25 0.29 0.28 0.3 0.27 0.15 0.15 0.16 0.17 0.17 0.2 0.2 0.17 0.18 0.16 0.2 0.2 0.18 0.18
29.79% TA 0.36 0.37 0.34 0.34 0.36 0.31 0.29 0.31 0.26 0.33 0.38 0.36 0.38 0.34 0.39 0.37 0.38 0.36 0.35 0.37 0.36 0.36 0.35 0.39 0.34 0.35 0.35 0.35 0.34 0.36 0.38 0.37
26.04% AC 0.24 0.24 0.33 0.28 0.32 0.37 0.43 0.37 0.41 0.28 0.2 0.21 0.2 0.23 0.21 0.2 0.22 0.21 0.28 0.3 0.37 0.35 0.27 0.23 0.22 0.24 0.23 0.22 0.24 0.22 0.2 0.23
29.88% CC 0.45 0.42 0.34 0.45 0.4 0.34 0.31 0.35 0.27 0.37 0.26 0.29 0.26 0.29 0.26 0.31 0.27 0.32 0.41 0.39 0.36 0.32 0.37 0.38 0.38 0.38 0.41 0.43 0.38 0.39 0.37 0.41
19.35% GC 0.1 0.11 0.13 0.082 0.096 0.097 0.094 0.1 0.1 0.11 0.27 0.25 0.28 0.21 0.27 0.24 0.26 0.21 0.078 0.11 0.089 0.099 0.12 0.15 0.15 0.13 0.12 0.12 0.14 0.15 0.15 0.13
24.72% TC 0.21 0.23 0.2 0.18 0.19 0.19 0.17 0.18 0.22 0.24 0.27 0.26 0.25 0.27 0.26 0.25 0.25 0.26 0.22 0.2 0.18 0.24 0.24 0.24 0.25 0.25 0.24 0.23 0.25 0.25 0.28 0.23
30.42% AG 0.28 0.31 0.39 0.28 0.3 0.32 0.37 0.44 0.41 0.3 0.26 0.27 0.27 0.28 0.27 0.27 0.27 0.27 0.41 0.31 0.36 0.45 0.27 0.27 0.3 0.3 0.3 0.24 0.31 0.29 0.27 0.25
11.84% CG 0.31 0.24 0.21 0.36 0.23 0.26 0.17 0.12 0.18 0.24 0.18 0.18 0.15 0.19 0.15 0.17 0.14 0.17 0.25 0.24 0.19 0.2 0.23 0.19 0.21 0.21 0.21 0.26 0.22 0.21 0.2 0.23
29.89% GG 0.21 0.26 0.23 0.19 0.24 0.22 0.25 0.22 0.25 0.22 0.3 0.29 0.32 0.29 0.31 0.3 0.31 0.31 0.14 0.22 0.23 0.17 0.24 0.29 0.26 0.24 0.27 0.26 0.25 0.25 0.27 0.26
27.86% TG 0.21 0.2 0.18 0.17 0.23 0.2 0.21 0.22 0.16 0.24 0.26 0.26 0.26 0.25 0.26 0.26 0.28 0.26 0.2 0.23 0.22 0.19 0.26 0.25 0.23 0.25 0.23 0.25 0.22 0.24 0.25 0.26
26.69% AU 0.23 0.2 0.3 0.27 0.26 0.36 0.46 0.35 0.64 0.22 0.22 0.21 0.21 0.21 0.23 0.19 0.21 0.19 0.24 0.23 0.23 0.27 0.2 0.22 0.2 0.2 0.2 0.2 0.21 0.21 0.2 0.22
24.25% CU 0.45 0.44 0.32 0.4 0.39 0.34 0.21 0.23 0.13 0.38 0.21 0.25 0.19 0.29 0.18 0.31 0.2 0.32 0.44 0.29 0.31 0.33 0.42 0.33 0.36 0.4 0.36 0.38 0.38 0.29 0.33 0.36
20.77% GU 0.14 0.18 0.21 0.2 0.16 0.17 0.19 0.18 0.16 0.19 0.33 0.32 0.36 0.28 0.37 0.27 0.36 0.29 0.13 0.22 0.14 0.14 0.17 0.2 0.2 0.16 0.2 0.2 0.19 0.21 0.21 0.22
28.30% TU 0.18 0.18 0.17 0.13 0.19 0.13 0.14 0.23 0.075 0.21 0.24 0.23 0.24 0.22 0.21 0.22 0.22 0.2 0.19 0.26 0.32 0.27 0.21 0.25 0.24 0.23 0.24 0.22 0.22 0.29 0.27 0.21
CD4⁺T-FS185
CD4⁺T-FS186
CD4⁺T-FS187
CD4⁺T-FS188
CD4⁺T-FS189
hESC-H9-FS197
hESC-H9-FS198
hESC-H9-FS199
hESC-H9-FS201
hESC-H9-FS303
DLTB-FS257
DLTB-FS259
DLTB-FS261
DLTB-FS263
TLTB-FS258
TLTB-FS260
TLTB-FS262
TLTB-FS264
WB-GTP control-FS195
WB-GTP PSTD-FS196
HCT116-FS193
HEK293T-FS203
HEK293T-FS305
HEK293T-FS326
RNH2A-KO T3-8-FS300
RNH2A-KO T3-8-FS306
RNH2A-KO T3-8-FS309
RNH2A-KO T3-8-FS327
RNH2A-KO T3-17-FS301
RNH2A-KO T3-17-FS307
RNH2A-KO T3-17-FS310
RNH2A-KO T3-17-FS329

### Figure S12

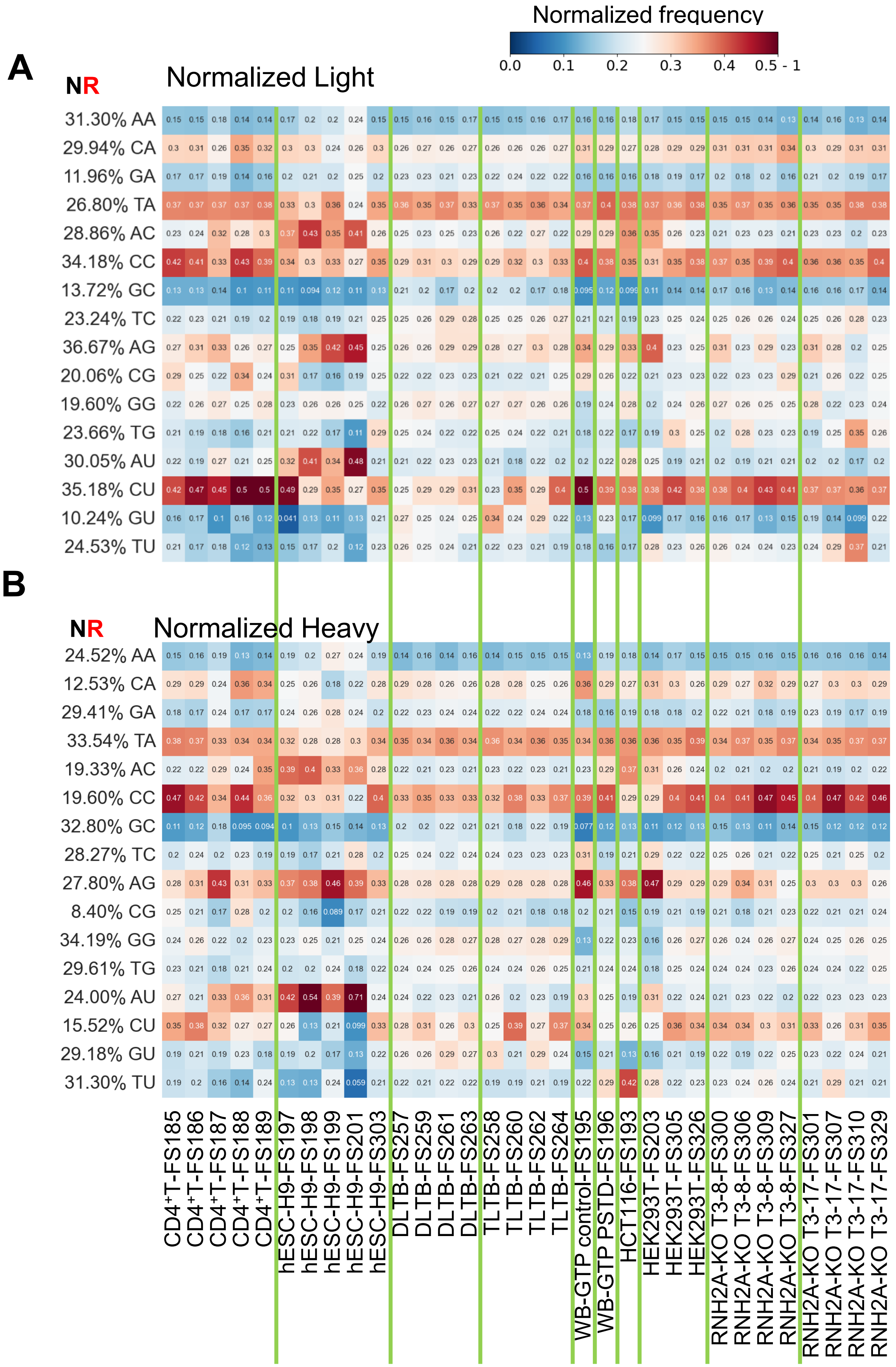
Normalized frequency
0.0
0.1
0.2
0.3
0.4
0.5 - 1
A
NR
Normalized Light
31.30% AA
29.94% CA
11.96% GA
26.80% TA
28.86% AC
34.18% CC
13.72% GC
23.24% TC
36.67% AG
20.06% CG
19.60% GG
23.66% TG
30.05% AU
35.18% CU
10.24% GU
24.53% TU
B
NR
Normalized Heavy
24.52% AA
12.53% CA
29.41% GA
33.54% TA
19.33% AC
19.60% CC
32.80% GC
28.27% TC
27.80% AG
8.40% CG
34.19% GG
29.61% TG
24.00% AU
15.52% CU
29.18% GU
31.30% TU
CD4+T-FS185
CD4+T-FS186
CD4+T-FS187
CD4+T-FS188
CD4+T-FS189
hESC-H9-FS197
hESC-H9-FS198
hESC-H9-FS199
hESC-H9-FS201
hESC-H9-FS303
DLTB-FS257
DLTB-FS259
DLTB-FS261
DLTB-FS263
TLTB-FS258
TLTB-FS260
TLTB-FS262
TLTB-FS264
WB-GTP control-FS195
WB-GTP PSTD-FS196
HCT116-FS193
HEK293T-FS203
HEK293T-FS305
HEK293T-FS326
RNH2A-KO T3-8-FS300
RNH2A-KO T3-8-FS306
RNH2A-KO T3-8-FS309
RNH2A-KO T3-8-FS327
RNH2A-KO T3-17-FS301
RNH2A-KO T3-17-FS307
RNH2A-KO T3-17-FS310
RNH2A-KO T3-17-FS329

### Figure S13

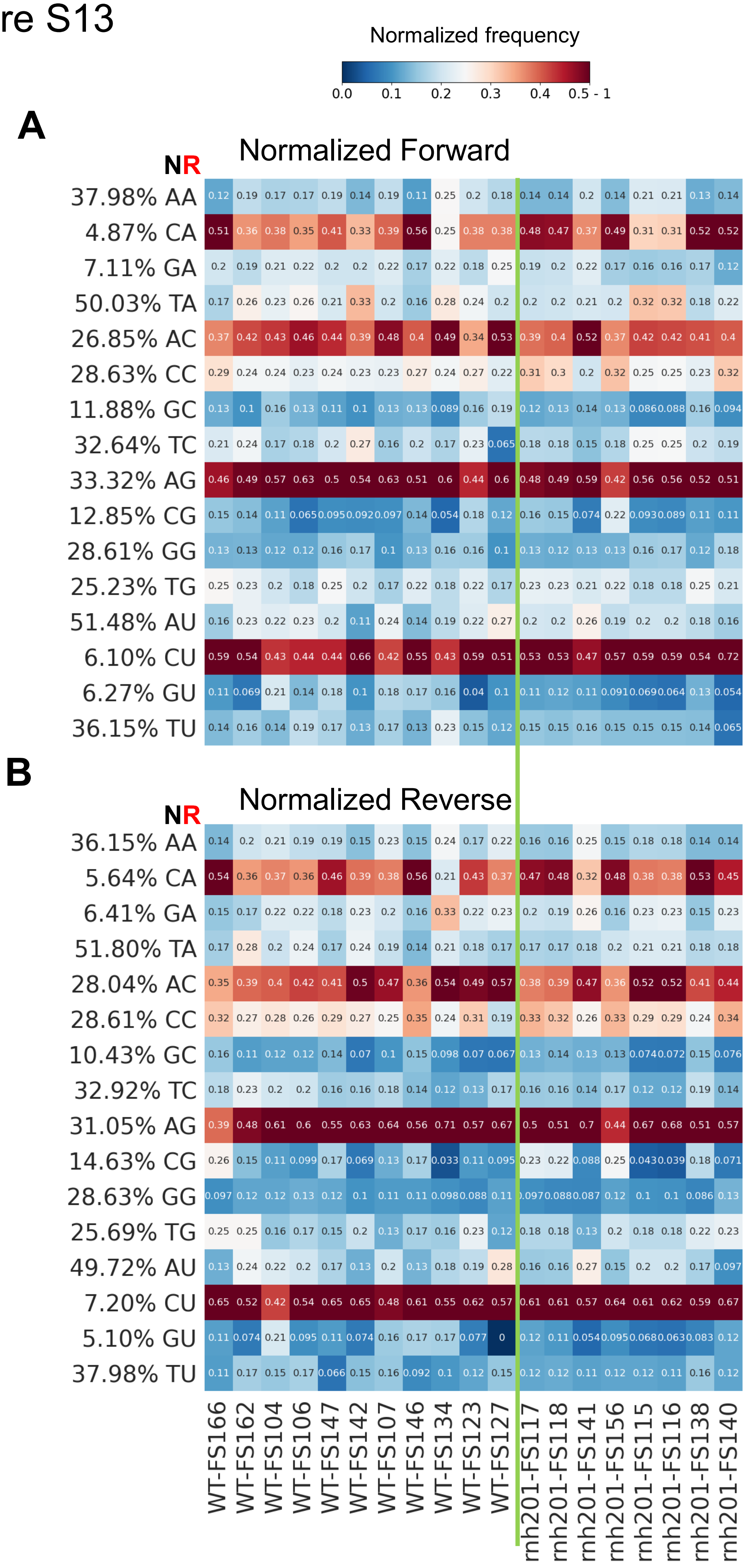
Normalized frequency
0.0 0.1 0.2 0.3 0.4 0.5 - 1
A
Normalized Forward
NR
37.98% AA 0.12 0.19 0.17 0.17 0.19 0.14 0.19 0.11 0.25 0.2 0.18 0.14 0.14 0.2 0.14 0.21 0.21 0.13 0.14
4.87% CA 0.51 0.36 0.38 0.35 0.41 0.33 0.39 0.56 0.25 0.38 0.38 0.48 0.47 0.37 0.49 0.31 0.31 0.52 0.52
7.11% GA 0.2 0.19 0.21 0.22 0.2 0.2 0.22 0.17 0.22 0.18 0.25 0.19 0.2 0.22 0.17 0.16 0.16 0.17 0.12
50.03% TA 0.17 0.26 0.23 0.26 0.21 0.33 0.2 0.16 0.28 0.24 0.2 0.2 0.2 0.21 0.2 0.32 0.32 0.18 0.22
26.85% AC 0.37 0.42 0.43 0.46 0.44 0.39 0.48 0.4 0.49 0.34 0.53 0.39 0.4 0.52 0.37 0.42 0.42 0.41 0.4
28.63% CC 0.29 0.24 0.24 0.23 0.24 0.23 0.23 0.27 0.24 0.27 0.22 0.31 0.3 0.2 0.32 0.25 0.25 0.23 0.32
11.88% GC 0.13 0.1 0.16 0.13 0.11 0.1 0.13 0.13 0.089 0.16 0.19 0.12 0.13 0.14 0.13 0.086 0.088 0.16 0.094
32.64% TC 0.21 0.24 0.17 0.18 0.2 0.27 0.16 0.2 0.17 0.23 0.065 0.18 0.18 0.15 0.18 0.25 0.25 0.2 0.19
33.32% AG 0.46 0.49 0.57 0.63 0.5 0.54 0.63 0.51 0.6 0.44 0.6 0.48 0.49 0.59 0.42 0.56 0.56 0.52 0.51
12.85% CG 0.15 0.14 0.11 0.065 0.095 0.092 0.097 0.14 0.054 0.18 0.12 0.16 0.15 0.074 0.22 0.093 0.089 0.11 0.11
28.61% GG 0.13 0.13 0.12 0.12 0.16 0.17 0.1 0.13 0.16 0.16 0.1 0.13 0.12 0.13 0.13 0.16 0.17 0.12 0.18
25.23% TG 0.25 0.23 0.2 0.18 0.25 0.2 0.17 0.22 0.18 0.22 0.17 0.23 0.23 0.21 0.22 0.18 0.18 0.25 0.21
51.48% AU 0.16 0.23 0.22 0.23 0.2 0.11 0.24 0.14 0.19 0.22 0.27 0.2 0.2 0.26 0.19 0.2 0.2 0.18 0.16
6.10% CU 0.59 0.54 0.43 0.44 0.44 0.66 0.42 0.55 0.43 0.59 0.51 0.53 0.53 0.47 0.57 0.59 0.59 0.54 0.72
6.27% GU 0.11 0.069 0.21 0.14 0.18 0.1 0.18 0.17 0.16 0.04 0.1 0.11 0.12 0.11 0.091 0.069 0.064 0.13 0.054
36.15% TU 0.14 0.16 0.14 0.19 0.17 0.13 0.17 0.13 0.23 0.15 0.12 0.15 0.15 0.16 0.15 0.15 0.15 0.14 0.065
B
Normalized Reverse
NR
36.15% AA 0.14 0.2 0.21 0.19 0.19 0.15 0.23 0.15 0.24 0.17 0.22 0.16 0.16 0.25 0.15 0.18 0.18 0.14 0.14
5.64% CA 0.54 0.36 0.37 0.36 0.46 0.39 0.38 0.56 0.21 0.43 0.37 0.47 0.48 0.32 0.48 0.38 0.38 0.53 0.45
6.41% GA 0.15 0.17 0.22 0.22 0.18 0.23 0.2 0.16 0.33 0.22 0.23 0.2 0.19 0.26 0.16 0.23 0.23 0.15 0.23
51.80% TA 0.17 0.28 0.2 0.24 0.17 0.24 0.19 0.14 0.21 0.18 0.17 0.17 0.17 0.18 0.2 0.21 0.21 0.18 0.18
28.04% AC 0.35 0.39 0.4 0.42 0.41 0.5 0.47 0.36 0.54 0.49 0.57 0.38 0.39 0.47 0.36 0.52 0.52 0.41 0.44
28.61% CC 0.32 0.27 0.28 0.26 0.29 0.27 0.25 0.35 0.24 0.31 0.19 0.33 0.32 0.26 0.33 0.29 0.29 0.24 0.34
10.43% GC 0.16 0.11 0.12 0.12 0.14 0.07 0.1 0.15 0.098 0.07 0.067 0.13 0.14 0.13 0.13 0.074 0.072 0.15 0.076
32.92% TC 0.18 0.23 0.2 0.2 0.16 0.16 0.18 0.14 0.12 0.13 0.17 0.16 0.16 0.14 0.17 0.12 0.12 0.19 0.14
31.05% AG 0.39 0.48 0.61 0.6 0.55 0.63 0.64 0.56 0.71 0.57 0.67 0.5 0.51 0.7 0.44 0.67 0.68 0.51 0.57
14.63% CG 0.26 0.15 0.11 0.099 0.17 0.069 0.13 0.17 0.033 0.11 0.095 0.23 0.22 0.088 0.25 0.043 0.039 0.18 0.071
28.63% GG 0.097 0.12 0.12 0.13 0.12 0.1 0.11 0.11 0.098 0.088 0.11 0.097 0.088 0.087 0.12 0.1 0.1 0.086 0.13
25.69% TG 0.25 0.25 0.16 0.17 0.15 0.2 0.13 0.17 0.16 0.23 0.12 0.18 0.18 0.13 0.2 0.18 0.18 0.22 0.23
49.72% AU 0.13 0.24 0.22 0.2 0.17 0.13 0.2 0.13 0.18 0.19 0.28 0.16 0.16 0.27 0.15 0.2 0.2 0.17 0.097
7.20% CU 0.65 0.52 0.42 0.54 0.65 0.65 0.48 0.61 0.55 0.62 0.57 0.61 0.61 0.57 0.64 0.61 0.62 0.59 0.67
5.10% GU 0.11 0.074 0.21 0.095 0.11 0.074 0.16 0.17 0.17 0.077 0 0.12 0.11 0.054 0.095 0.068 0.063 0.083 0.12
37.98% TU 0.11 0.17 0.15 0.17 0.066 0.15 0.16 0.092 0.1 0.12 0.15 0.12 0.12 0.11 0.12 0.12 0.11 0.16 0.12
WT-FS166
WT-FS162
WT-FS104
WT-FS106
WT-FS147
WT-FS142
WT-FS107
WT-FS146
WT-FS134
WT-FS123
WT-FS127
rnh201-FS117
rnh201-FS118
rnh201-FS141
rnh201-FS156
rnh201-FS115
rnh201-FS116
rnh201-FS138
rnh201-FS140

### Figure S14

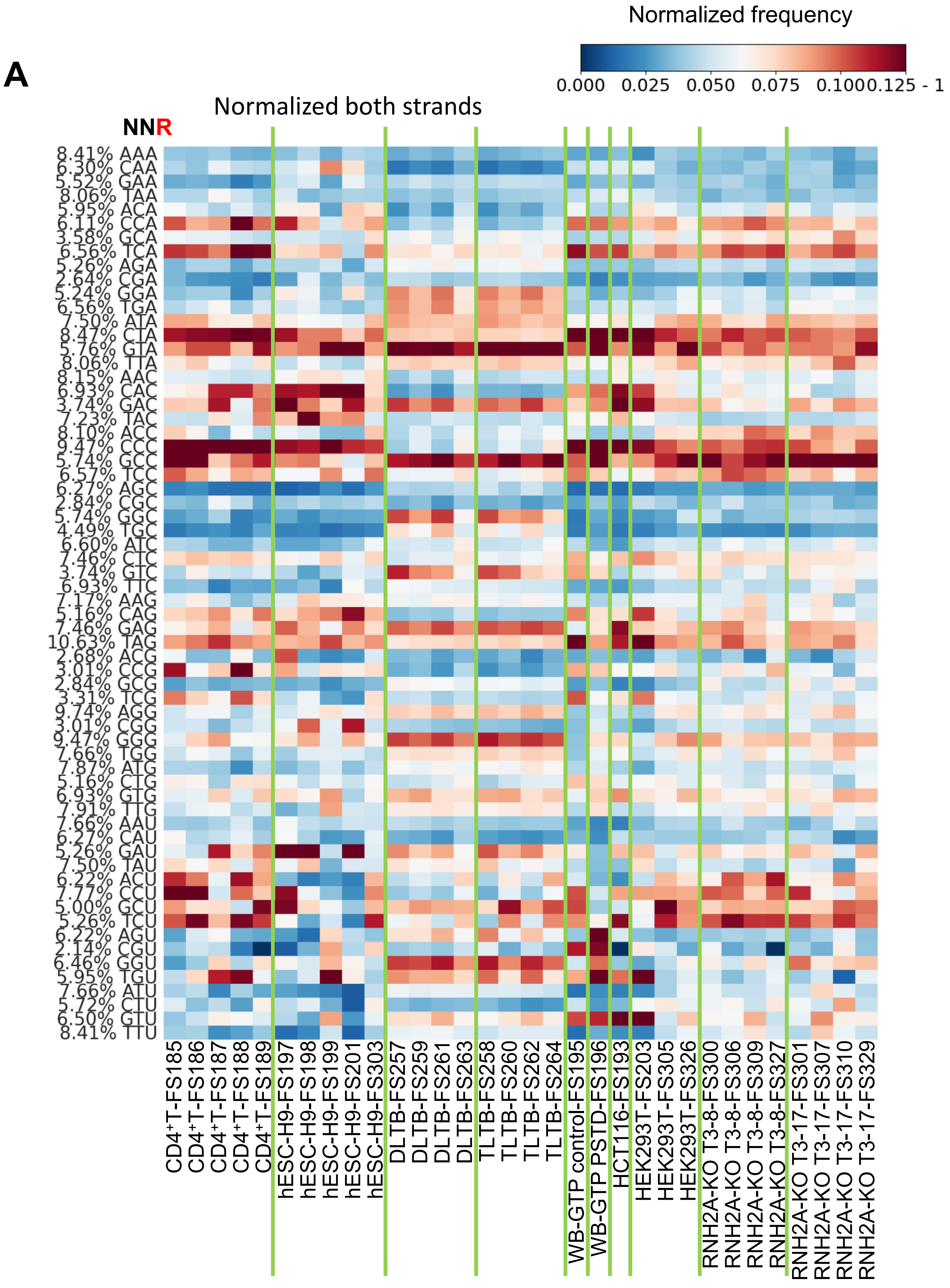
A
Normalized frequency
0.000
0.025
0.050
0.075
0.1000.125 - 1
Normalized both strands
NNR
8.41% AAA
6.30% CAA
5.52% GAA
8.06% TAA
5.95% ACA
6.11% CCA
3.58% GCA
6.56% TCA
5.26% AGA
2.64% CGA
5.24% GGA
6.56% TGA
7.50% ATA
8.47% CTA
5.76% GTA
8.06% TTA
8.15% AAC
6.93% CAC
3.74% GAC
7.23% TAC
8.10% ACC
9.47% CCC
5.74% GCC
6.57% TCC
6.27% AGC
2.84% CGC
5.74% GGC
4.49% TGC
6.60% ATC
7.46% CTC
3.74% GTC
6.93% TTC
7.17% AAG
5.16% CAG
7.46% GAG
10.63% TAG
2.68% ACG
3.01% CCG
2.84% GCG
3.31% TCG
9.74% AGG
3.01% CGG
9.47% GGG
7.66% TGG
7.87% ATG
5.16% CTG
6.93% GTG
7.91% TTG
7.66% AAU
6.27% CAU
5.26% GAU
7.50% TAU
6.22% ACU
7.77% CCU
5.00% GCU
5.26% TCU
6.22% AGU
2.14% CGU
6.46% GGU
5.95% TGU
7.66% ATU
5.72% CTU
6.50% GTU
8.41% TTU
CD4+T-FS185
CD4+T-FS186
CD4+T-FS187
CD4+T-FS188
CD4+T-FS189
hESC-H9-FS197
hESC-H9-FS198
hESC-H9-FS199
hESC-H9-FS201
hESC-H9-FS303
DLTB-FS257
DLTB-FS259
DLTB-FS261
DLTB-FS263
TLTB-FS258
TLTB-FS260
TLTB-FS262
TLTB-FS264
WB-GTP control-FS195
WB-GTP PSTD-FS196
HCT116-FS193
HEK293T-FS203
HEK293T-FS305
HEK293T-FS326
RNH2A-KO T3-8-FS300
RNH2A-KO T3-8-FS306
RNH2A-KO T3-8-FS309
RNH2A-KO T3-8-FS327
RNH2A-KO T3-17-FS301
RNH2A-KO T3-17-FS307
RNH2A-KO T3-17-FS310
RNH2A-KO T3-17-FS329

### Figure S14 continued

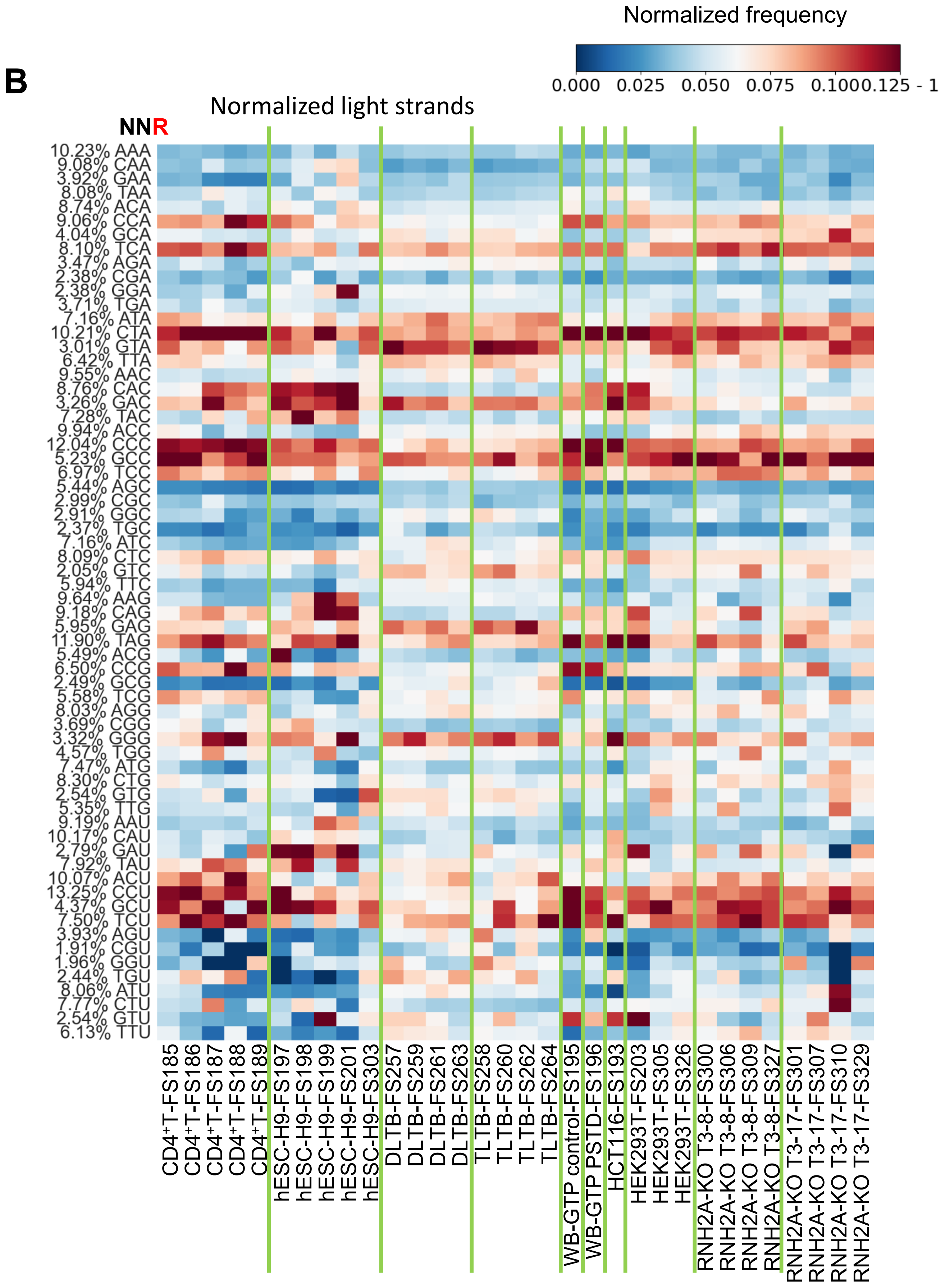

Figure S14 continued

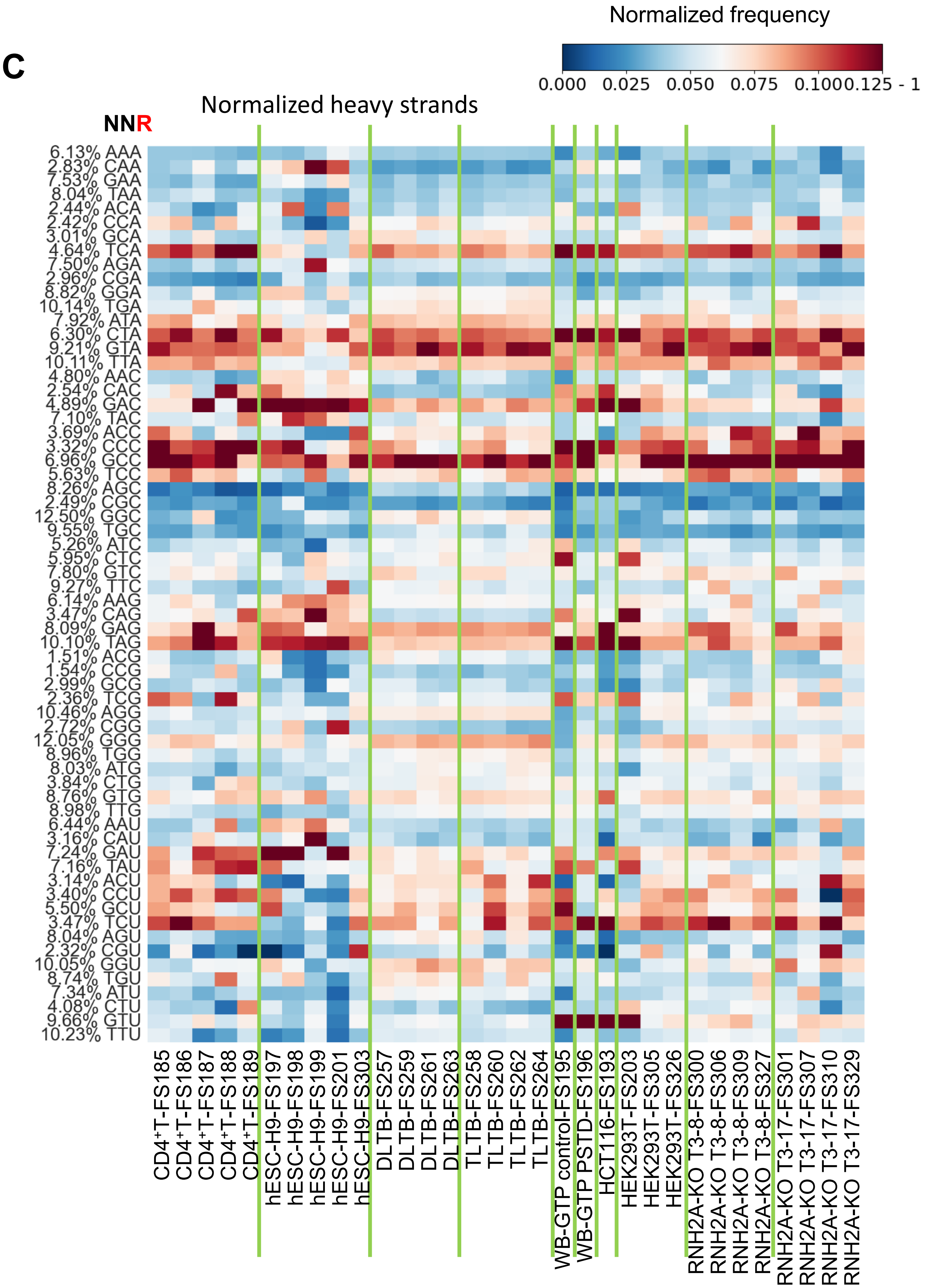

Figure S15

A

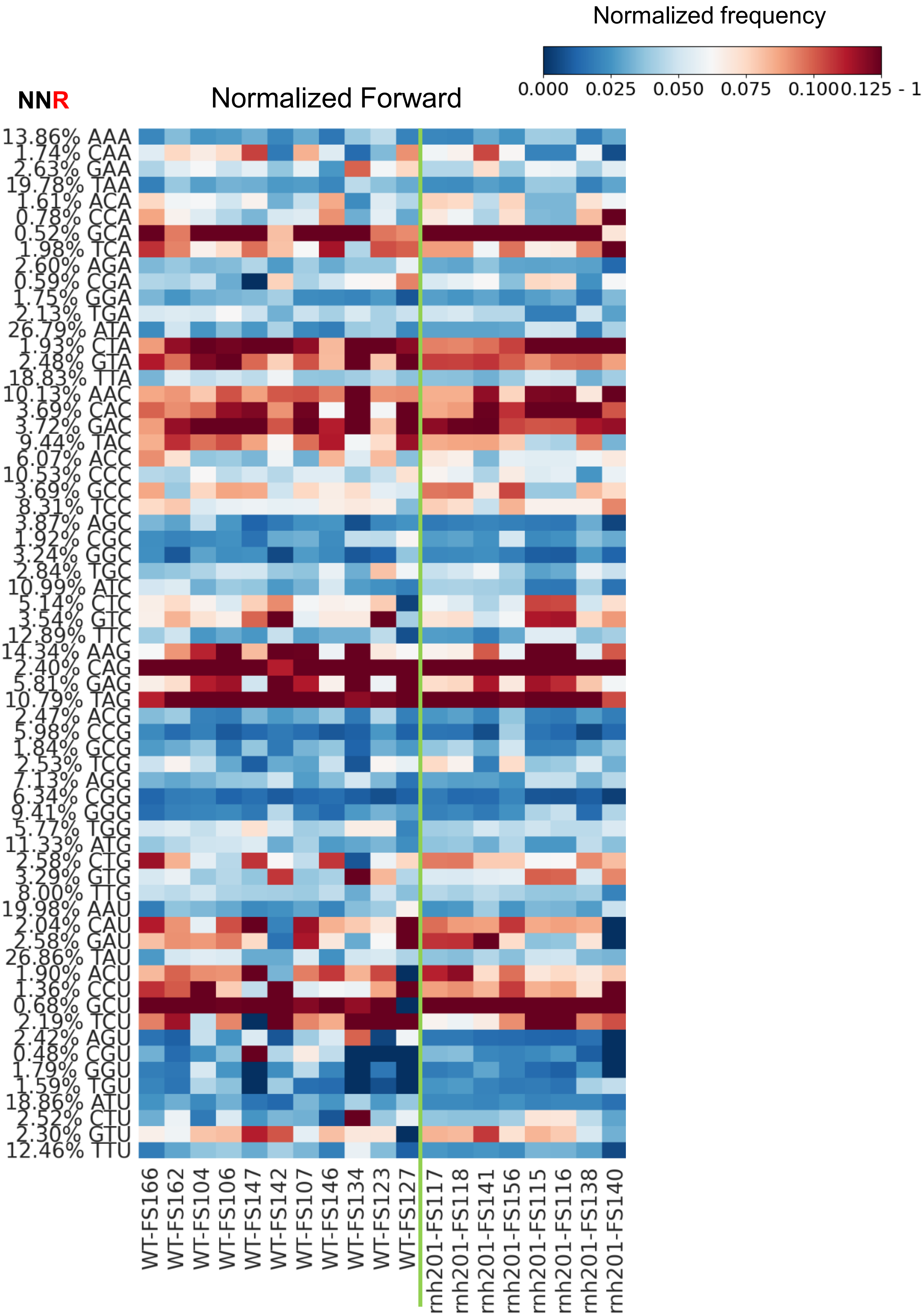
Normalized frequency
0.000 0.025 0.050 0.075 0.1000.125 - 1
NNR
Normalized Forward
13.86% AAA
1.74% CAA
2.63% GAA
19.78% TAA
1.61% ACA
0.78% CCA
0.52% GCA
1.98% TCA
2.60% AGA
0.59% CGA
1.75% GGA
2.13% TGA
26.79% ATA
1.93% CTA
2.48% GTA
18.83% TTA
10.13% AAC
3.69% CAC
3.72% GAC
9.44% TAC
6.07% ACC
10.53% CCC
3.69% GCC
8.31% TCC
3.87% AGC
1.92% CGC
3.24% GGC
2.84% TGC
10.99% ATC
5.14% CTC
3.54% GTC
12.89% TTC
14.34% AAG
2.40% CAG
5.81% GAG
10.79% TAG
2.47% ACG
5.98% CCG
1.84% GCG
2.53% TCG
7.13% AGG
6.34% CGG
9.41% GGG
5.77% TGG
11.33% ATG
2.58% CTG
3.29% GTG
8.00% TTG
19.98% AAU
2.04% CAU
2.58% GAU
26.86% TAU
1.90% ACU
1.36% CCU
0.68% GCU
2.19% TCU
2.42% AGU
0.48% CGU
1.79% GGU
1.59% TGU
18.86% ATU
2.52% CTU
2.30% GTU
12.46% TTU
WT-FS166
WT-FS162
WT-FS104
WT-FS106
WT-FS147
WT-FS142
WT-FS107
WT-FS146
WT-FS134
WT-FS123
WT-FS127
rnh201-FS117
rnh201-FS118
rnh201-FS141
rnh201-FS156
rnh201-FS115
rnh201-FS116
rnh201-FS138
rnh201-FS140

Figure S15 continued

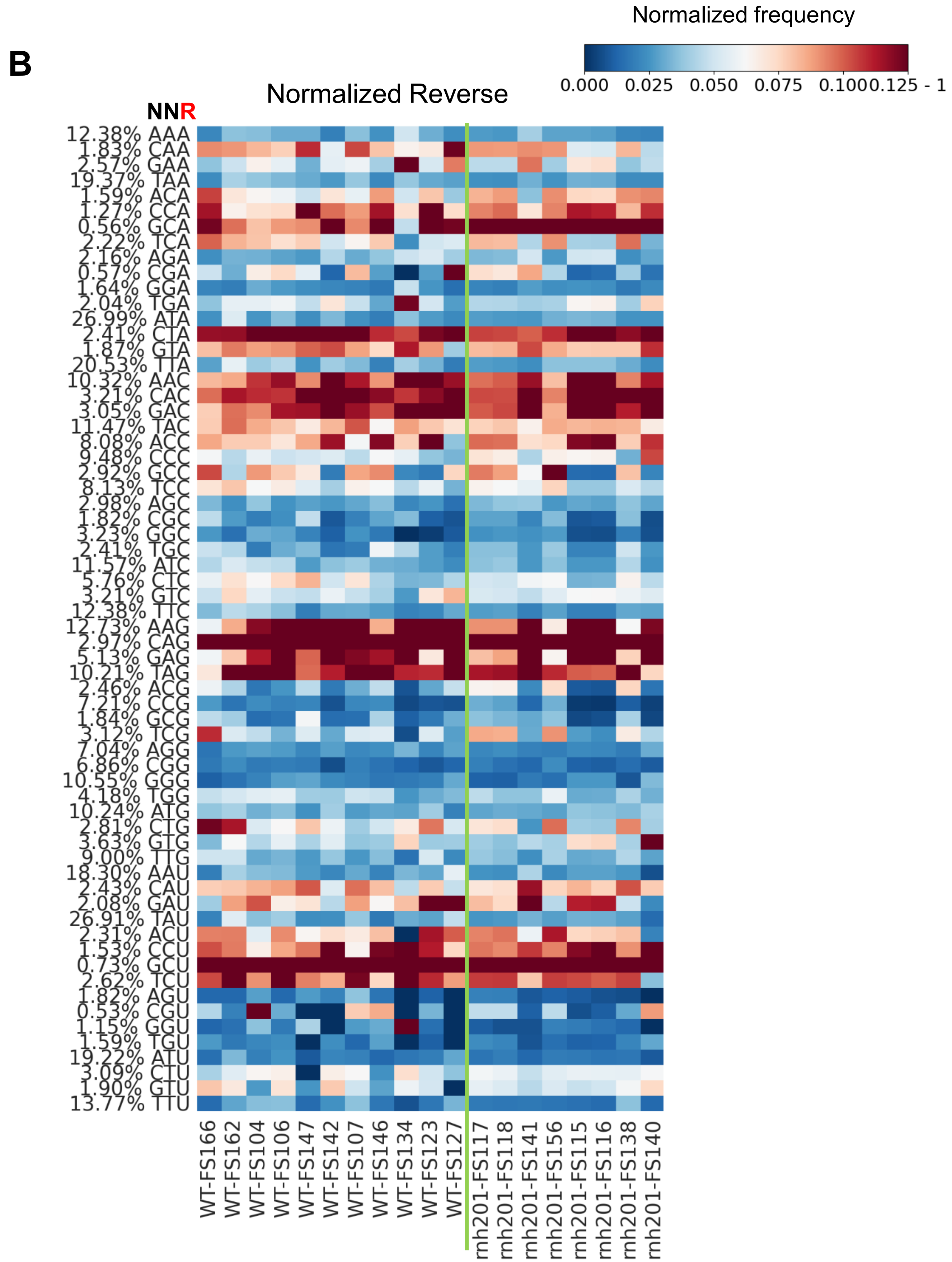
